# Coloring Single Nanoparticle Trajectory in Live Cell with its Own History: a Presuppositionless Preprocessing Approach

**DOI:** 10.1101/580480

**Authors:** Hansen Zhao, Zhenrong Huang, Feng Ge, Xiangjun Shi, Bin Xiong, Xuebin Liao, Zonghua Wang, Sichun Zhang, Xinrong Zhang, Yan He

## Abstract

Analyzing single particle trajectories is a prominent issue in understanding complex dynamics such as nanoparticle-cell interactions. Existing methods treat data points as isolated “atoms” and use predefined mechanical models to “frame” their complicated relationship. Herein, we propose a “historical evolution” based model-free strategy. It allows spatiotemporal heterogeneity embedded in a trajectory to self-emerge as consecutive colored segments before any model assumption, provide both an overall picture and local state transitions on the particle movement with minimum information loss, and inspire further model-based investigation. We demonstrate with simulations and experiments that the underlying mechanisms of various time-series and motion states of single nanoparticles on live cell membranes could all be revealed successfully. Since complexity studies at different levels of molecules, particles, cells, human beings, vehicles, and even stars could all be reduced to analyzing spatiotemporal trajectories of “single particles”, this presuppositionless approach will help fundamental researches on many important systems.

**Impact Statement:** A preprocessing strategy for single particle trajectory analysis is established by providing an intuitive global pattern from “historical experiences” of the particle without predefining any mechanical models.

## 1. Introduction

Investigation of the interactions between nanoparticles (NPs) and the live cell are crucial in the understanding of important biological processes such as nanoparticle cytotoxicity (1), drug delivery (2), gene therapy (3) and cellular endocytosis (4). Toward this end, tracking and analyzing the motions of individual nanoparticles in situ in real time, which could provide the spatial and temporal distribution of heterogeneity associated with the complex environments on cell surface, has attracted continuous interest (5–9). Taking advantages of rapid developments of optical imaging methods, researchers have been able to obtain the spatiotemporal series of multiple feature parameters such as the fluorescence and scattering intensities, the spectra and colors (10), and the rotation and orientation angles of individual NPs (9, 11, 12). However, the efforts to extract meaningful information and uncover the complex physical and biological mechanisms are always entangled with the stochastic natures of the NP motion, transient NP interaction with cell surface (13, 14), and varying distribution of surface micro-domains as well as the detection error.

Given the high spatiotemporal complexity of NP-cell interactions, the primary issue is to find an appropriate approach to differentiate and describe the particle movement. As a special time-series, a single particle trajectory is the consequence of a free particle exploring an unknown environment, and has several unique characteristics. 1) Each data point results from an instantaneous reaction with the local environment and carries certain physical meaning. 2) The reactive property of the environment has certain degree of continuity, which has scales greater than the spatial and temporal resolution of the microscopic measurement and could be designated as a local state. 3) There is an overall picture on the different environmental states and their relationship embedded in the particle trajectory within limited spatiotemporal range.

To begin with, conventional workflow must predefine the local environments and external forces that surround and restrict the single NPs, i.e. the motion-states the NPs could exhibit, and stipulate a framework based on *a priori* physical models. The data for analysis are often generated from the particle trajectory using subsequence or moving window operation. The general assumption is that during the NP-cell interaction process, the NP motion-state within a short time window is stable or statistically predictable, and different NP states are independent and limited in number (15). These underlying states could be recovered by either numerical fitting or by machine learning based model assignment, where the sub-trajectories are treated as atomistic fragment or event and are generally considered to be isolated from the adjacent ones. For example, by assuming that the mean-square-displacement in a subsequence can be described by a general equation of the anomalous diffusion, different particle states can be separated based on empirically determined thresholds of diffusion coefficients and asymmetry factors (16). Alternatively, by assuming the whole trajectory being a Markov process containing multiple states, and pre-estimating the initial state probability, emission probability and state transition probability, the hidden Markov model (HMM) method could be used to evaluate the sequence of the hidden states step-by-step and infer the underlying mechanism of particle motion (17, 18). To relax the prerequisite on either numerical or statistical description of the transient dynamics, some people resort to supervised clustering. Using empirically selected or simulated sub-trajectories to feed the classifier in advance, support vector machine (19), random forest (20) and Bayesian inference (21) have been proved to be reliable for dynamic state sequence differentiation. All these reductive strategies are largely applicable to reveal the general characteristics of the system embedded in the NP trajectories, but the assumptions on *a priori* states or models often suffer from oversimplification of the NP-cell interactions that are intrinsically heterogeneous and dynamic in time and space, especially for studies on rarely explored systems.

A possible way to address this issue is to transform the complicated dynamic process of individuals over time into a more intuitive picture with minimal information losses before further model definition and investigation. To achieve this goal, we have to establish a presuppositionless model-free strategy that have the internal states or inner differences emerge on their own. While this possibility has not been considered seriously in physical-model based fundamental research, the corresponding issue in philosophy, namely how to “return to things themselves”, has been examined in-depth for centuries by European philosophers during the study of phenomenology of time-consciousness (22). The core issue is that how a temporal object consisting of successive moments appears as the temporal unity of multiple consecutive states (Figure supplement 1). For example, a person is listening to a melody. How can he perceive detailed changes of the music while enjoying the overall beauty of the melody? Obviously, these perceived details in sound are arranged sequentially over time, but neither are monotonous notes nor mechanical combinations of notes, and cannot be pre-communicated with “musical models”. According to the phenomenology studies, at each time point, the basic unit of perception is not an atomistic moment irrelevant to others, but a conscious experience-fragment with some temporal depth anchored by the current point. Each experience fragment is composed of a certain number of consecutive moments before that point, so the successive experiences are different, but are interrelated due to their overlap in time. Such “succession of experiences” is the foundation by which the temporal change can be perceived. However, without a predefined *a priori* criterion, just a sequence of experiences of isolated short musical fragments are not sufficient to produce the perceived musical unity as well as the detailed states on the melody. Hence, a special operation at a higher conscious level is implemented to differentiate the short experience-fragments from each other, and simultaneously coalesce them in the temporal order, so that the person can actually “experience the succession” or perceive the temporality of the melody, i.e. the past, present and future phases of the temporal object. Since an experience-fragment is just a short historical fragment, the whole process is essentially a kind of “comparative historical analysis”. The theoretical basis with more philosophical arguments are provided in the Appendix 1 and 2. Nevertheless, this concept has remained untouched in natural scientific research due to the difficulty in making it into an objective and reproducible computational framework.

The coming of Data Era makes it possible to reconcile the physical science and phenomenology philosophy at the data science level. (23–25). Data are acquired objectively, but are little meaningful without being subjectively interpreted or visualized appropriately. If the data is highly fragmented or the amount of data is very limited, people would have to resort to some predefined prototypes to rationalize the otherwise isolated, seemingly clueless data points or fragments. With the size of available data growing fast exponentially, it becomes possible to “have the data speak for themselves” through data-driven approaches (26, 27). Inspired by the phenomenology study, in this work we propose a new model-free strategy to preprocess the single NP trajectory and uncover its overall spatiotemporal structure before further application of model-based analysis. We treat a trajectory as if it were a nanoscale melody having been experienced by the particle. To reveal the detailed states of the particle motion as well as their change over time as a unity, we conceptualize a special data pre-processing operator composed of 3 steps (Figure 1). 1) Construct a succession of overlapping experience-fragments of the particle with a specific predisposition. 2) Perform internal comparison or unsupervised clustering of all the experience-fragments to find their relative differences. 3) Color the trajectory accordingly to generate the perceivable experience of succession through segmentation of the particle motion. The resulting trajectory with multiple color segments can be further analyzed to extract more specific information based on the existing knowledge or models. Since there is a transformation from the “succession of experience” to the “experience of succession”, we designate the method by the acronym SEES. We demonstrate with simulated and experimentally obtained NP trajectories that, after applying the nodel-free SEES operator, the spatiotemporal dynamic states of NP-cell interactions could be successfully unveiled. This generalized data-phenomenology framework could give an overview of the complex spatiotemporal particle dynamics before the conventional model definition and promote the application of data-driven techniques in complexity studies.

**Figure 1.**
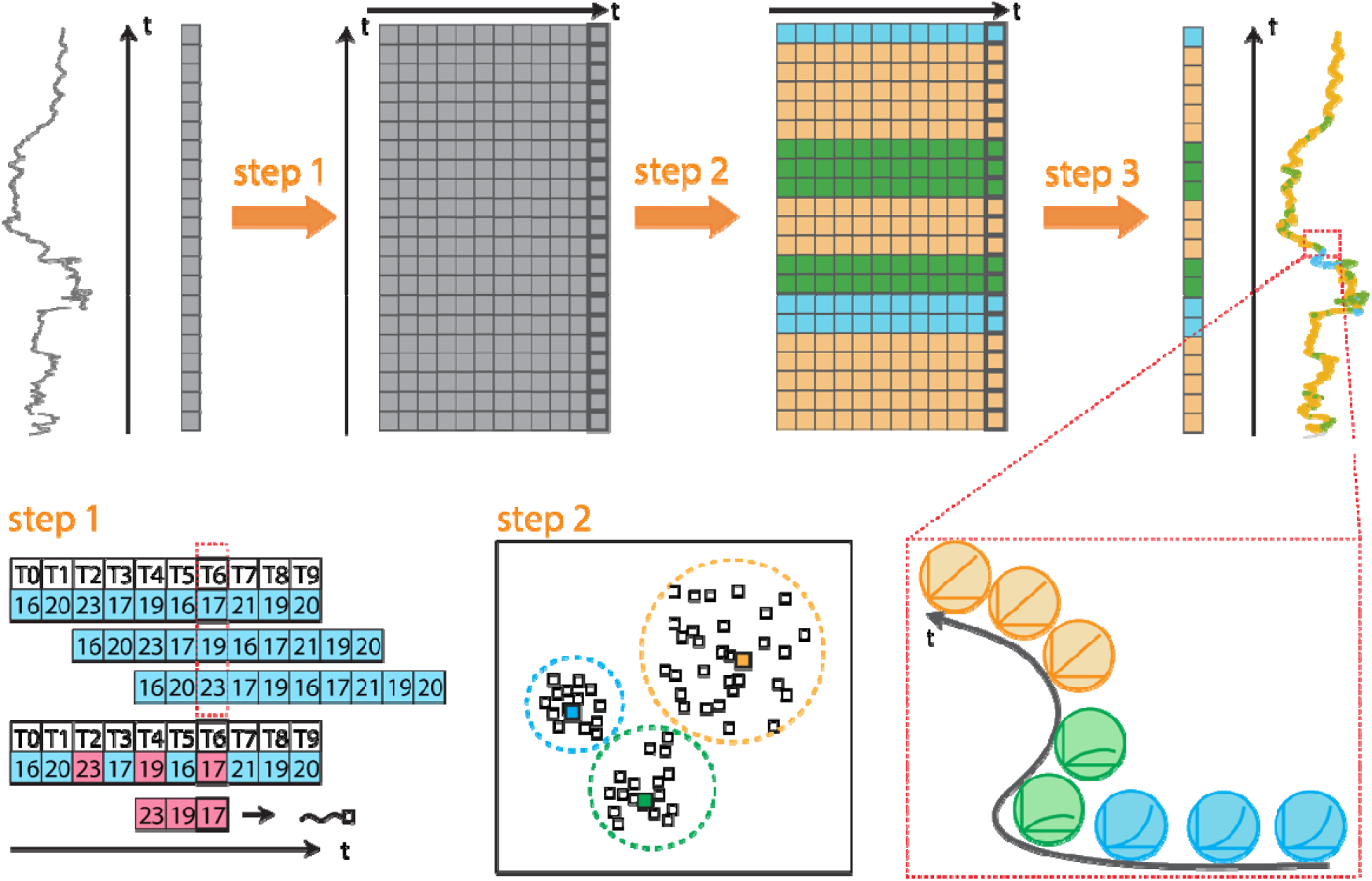
The 3-step SEES operator: step 1, construct the sequence of historical experience-fragments by the time-lag operation; step 2, apply k-means unsupervised clustering to differentiate the experience-vectors in high dimensional space; step 3, map the color tags of the clustering results back to the trajectory. The color fragments formed can then be analyzed by using existing physical models.

## 2. Description of the SEES Method

The first step of the SEES operator is to construct a succession of short experiences out of the single particle trajectory. The key here is not to view a trajectory obtained from a complex system as a simple assembly of separated atomistic events bound by fixed external laws or mechanical states, but as a consecutive sequence of cross-related historical experiences anchored by their current points. Each current-point carries an experience-fragment composed of its past points, and itself is also the immediate past point in the experience-fragment of the following point, and so on. This step is technically similar to the sliding window operation but is conceptually different. The later only intends to reduce noise from the data, and still treats each window as an isolated atomic fragment that can be collapsed into either a fixed value by averaging or a predefined kinetic phase by fitting, which inevitably suffers from information loss. The former treats each fragment as a temporal vector that keeps the chronological order of the data points, in which all the dynamic information is preserved. We note that while building the sequence of experience-vectors does not need predefined *a priori* models, it cannot be done without certain “intention” that is based on the prior knowledge of the system. Just as there are many perspectives to looking into the history of an individual, in what aspect the researcher intends to investigate determines how to generate the experience-vectors. For a single NP trajectory, we can focus on different features of the particle such as its velocity or orientation angle values, its rotational diffusion rate, and its spatial confinement, etc., and construct the experience-vector sequence accordingly. Since the purpose is to differ them internally, the historical length of the vectors is not crucial as long as it reaches a minimum. To select the appropriate value, well-developed non-linear time series analysis methods such as the Cao method (28) and the mutual information method (29) can be applied to provide a meaningful initial estimation, though the NP-cell interactions may not be chaotic.

The second step of the operator is to find the inner difference within the sequence of the NP experience-vectors by comparing each vector with all others simultaneously. This can be done via a number of machine-learning based techniques. Here, we chose the basic unsupervised k-means clustering method just to prove the concept. With the whole set of historical experience-vectors as the input, the output is the high-dimensional vector space being partitioned into a few categories with their cluster center. The criteria or metrics for the partitioning can be set based on the construction of the historical experience-vectors and the feature to be probed. The appropriate number of categories can be estimated according to the literature (30). We note that such action is similar in certain extent to the subsequence time series clustering, which has been reported to be inefficient in revealing the true physical structure of some types of time series (31). However, instead of focusing on the physical importance of the cluster centers, what the SEES operator does at this step is just to have their differences emerge but not necessarily to justify their physical nature. It is because that as internal comparison without *a priori* criteria, the differences between cluster centers are only relative and the classification of each experience-vector is probabilistic. To make the clustering results physically meaningful, their difference must be mapped back to the original spatiotemporal trajectory of the particle, and that leads to the next step of the operator.

The third step is to color the single NP trajectory according to the clustering difference. By assigning each cluster of experience-vectors a color and labeling every point on the trajectory using the color of the vector it carries, we obtain a trajectory labelled with multiple colors. Because of the existence of both local consistency and global difference, the same color points on the trajectory are usually not randomly distributed; they tend to assemble locally into multiple color segments, resulting in the emergence of a collective pattern of color continuities. The visually perceivable “experience of the succession” within each color segment could be attributed to the NP continuously experiences similar motion patterns or surrounding environments in time and space, and can be specified as a local state with a non-zero temporal depth. Hence the sequence of color change exhibits the variations and dynamic transitions of the local states. The physical meaning and characteristics of each state could be further analyzed according to previous knowledge on the process. We note that although the algorithm ensures it reproducible, the classification of an experience-vector is not deterministic but probabilistic, thus the temporal length and the boundaries of a color segment or the appearance of a local state are also probabilistic. However, such uncertainties are not stochastic and are linked globally by the wholeness of the trajectory, and the nondeterministic picture is consistent with the complex nature of NP-cell interactions, as long as a general picture describing the spatiotemporal distribution and transition of different local states could be obtained. Since the coloring is not just a result visualization, but important to subsequent model-based investigation, a relatively stable pattern resistant to slight parameter change is essential. This is usually achieved by gradually increase the length of the historical experience-vector to certain extent. In summary, the 3 steps of SEES operator are computationally and logically linked together to let the inner differences contained in the trajectory emerge on their own before model definitions, which could help to study those little explored systems. The implementations of the SEES operator, including constructions of the experience-vectors from a NP trajectory and the partition metrics in the k-means clustering algorithm, are provided in the Appendix 1 Note S2.

## 3. Method Evaluation

The implementations of the SEES operator, including constructions of the experience-vectors from a NP trajectory and the differentiation metrics in the k-means clustering algorithm, are provided in the Appendix 1 Note S2. Briefly, 3 ways of experience vector construction can be selected: univariate (U), mean squared displacement (MSD) and autocorrelation curve. The choice of the vector construction depends on which perspective of the spatiotemporal series data people want to focus. Generally, U, MSD and AC reflect the magnitude/tendency, dynamic confinement and self-similarity of the vector, respectively. As metrics for vector comparison, we used either Euclidean distance (ED) or cross-correlation (CC), which are sensitive to their magnitude and the variation trend, respectively.

In order to validate the method before using it to process the experimental data, we applied the SEES operator to simulate spatiotemporal series data with known ground truths (Appendix 1 Note S3). To examine the robustness and universal applicability of the operator, we generated 3 cases with totally different underlying physical mechanisms. The first example is a nanorod both rotating and revolving around an external axis simultaneously (Figure 2). The time-series obtained is the projection of its trajectory on one dimension, exhibiting both a local and a global periodic sinusoidal pattern. Targeting the tendency of data variation, we performed the SEES/U-CC operation as the univariate construction focuses the one-dimension series itself and the cross-correlation metric is sensitive to vector tendency. The resulting cluster centers indicate that the experience-vectors of the nanorod are correctly partitioned into two groups of ascending and descending tendency without physical or statistical presumption, and the periodical variation of the trends are rightly captured by the spontaneous accretion of the same color points on the colored trajectory.

**Figure 2.**
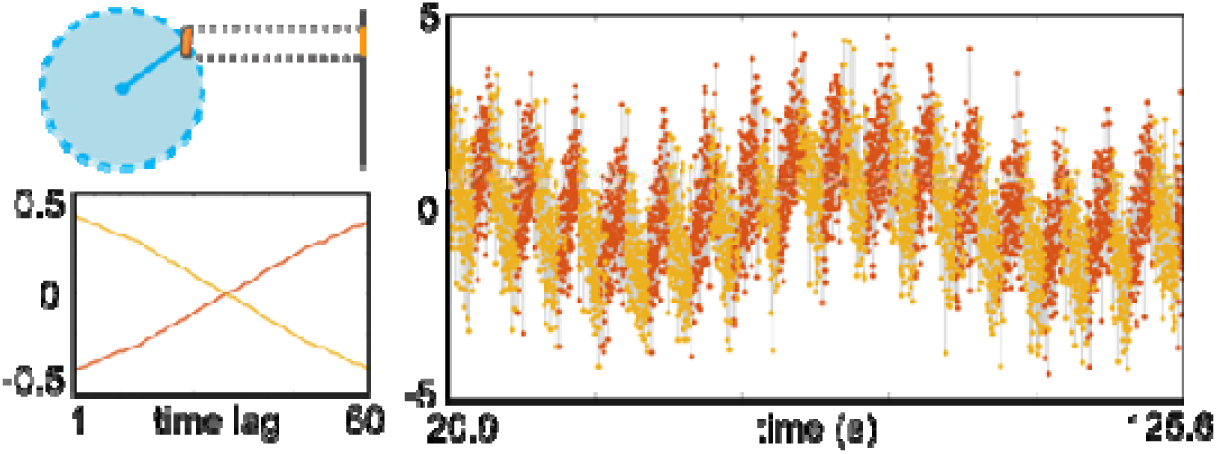
Nanorod rotation/revolution provides sinusoidal observations. The SEES/U-CC operation successfully revealed the periodic pattern and differentiated the ascending and descending tendency.

In the second case, we simulated a trajectory of a NP diffusing on a 1D membrane, whose moving rate varies at different regions due to the heterogeneity of the local environment (Figure 3). We applied the SEES/U-ED operator to the time series of NP velocity, in which the ED metric revealed the differences of the velocity magnitude that is related to the local diffusion coefficient. With the historical experience-vector length increasing from d=1 to 30, we can see how the underlying temporality of the NP motion emerges from the time-series gradually. When d is small, no clear pattern appears due to the stochastic nature of the transient NP rates. Increasing the vector length to 10 leads to a much better estimation of the transition of the diffusion states but still contained confusions. As d=30, the mean values of the cluster centers accurately match the ground truth. The resulting colored trajectory and spatiotemporal distribution of the color fragments are consistent with the original simulated one. The vector of the cluster center in each category shows approximately a flat curve whose mean value is comparable to the theoretical calculation according to the simulation. Surprisingly, not only the long regions but also the short regions of NP velocity variation are successfully recovered, though the boundaries and a few details are not precise. Close examination of the simulations with various NP diffusion coefficients and region sizes as input (Figure supplement 2) indicate that such good match could be attributed to the advantages of the large data set and the huge number of vector comparisons performed in the high dimensional space according to the Law of Large Number. Most of the physical models on single particle movements used nowadays were developed at times when the data acquisition and processing techniques were limited. Therefore, it is the data size that makes the presuppositionless SEES operator applicable.

**Figure 3.**
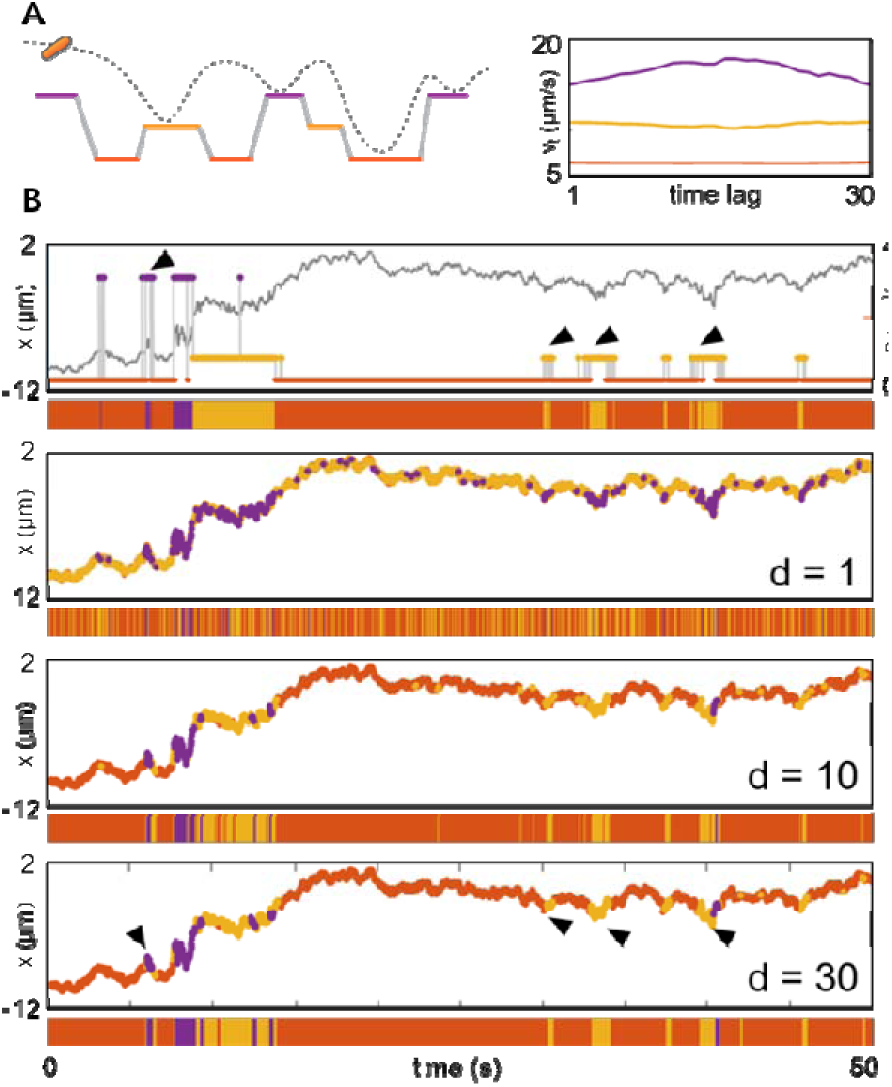
(A) Nanoparticle diffusion on a spatial heterogeneous 1D membrane. The schematic diagram of the model (left panel) and the cluster centers of the SEES/U-ED operation with vector length of 30. (B) The ground truth of the time series of NP velocity (row 1) and the results processed using the SEES/U-ED operator with the length of experience-vector, d, being set at d=1 (row 2), d=10 (row 3) and d=30 (row 4). When d=30, the clustering centers show high consistency to the expected mean velocities.

In the third example, we simulated a NP diffusing on a 2D cell membrane where a number of binding regions with different affinities distributed randomly. The NP jumps into or out of the confined regions randomly and undergoes confined or directed motions, respectively, producing a continuous trajectory (Figure 4). The possibility of state switching is not static throughout the process and does not depend on the immediately preceding state, but rather on the spatial location of the NP, and more specifically, the binding affinity of a particular region (Appendix 1 Note S3). Hence the state transition probability deviates from the basic assumption of HMM, but it has a clear biological significance: NPs coated with specific antibodies are more likely to bind to their corresponding antigens than other membrane proteins, and it is more difficult to escape from immune binding.

**Figure 4.**
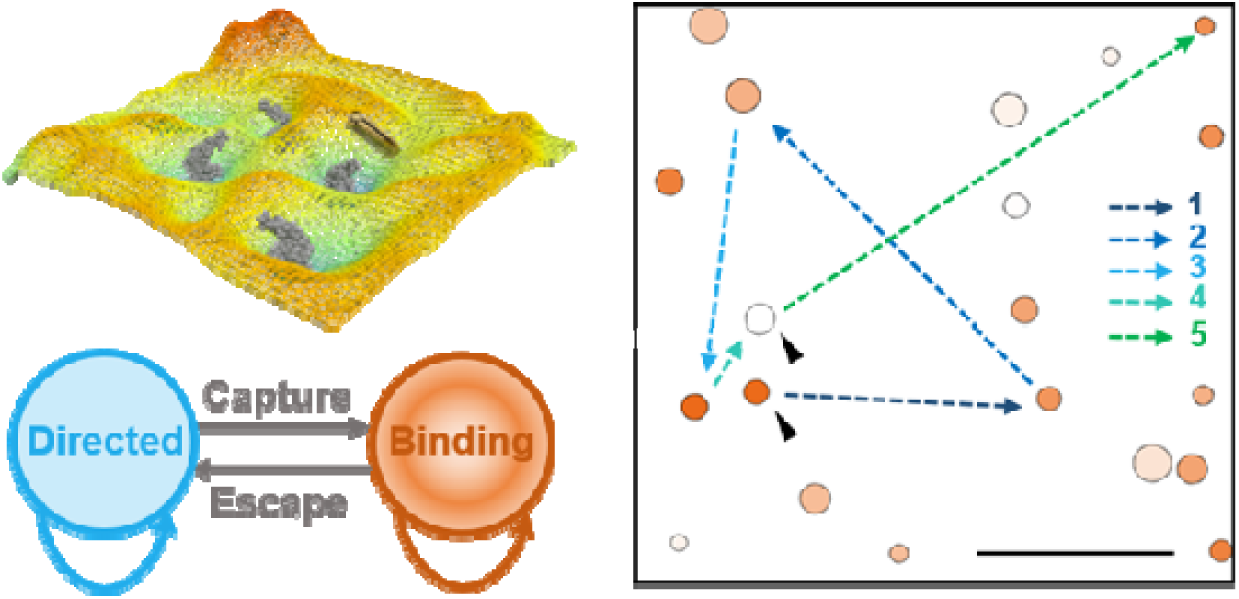
Binding/dissociation model simulates a nanoparticle interacting with a 2D cell membrane. Nanoparticles can switch between binding state and directed motion (dissociated) state depending on the spatial position. Binding sites with different local properties spread randomly on the surface are circled by orange color of different ligthness. The dash line arrows show the jumping trace of the nanoparticle from 10 s to 60 s of the simulation.

However, we could handle this situation by using two model-free SEES operators simultaneously. The SEES/MSD-ED operation on the NP position is sensitive to the historical confinement of the particle and indeed reveals the binding regions where the NP exploited. Meanwhile, the SEES/U-ED operation on the NP velocity series is more sensitive to the local diffusion coefficient inside the binding region (Figure 5A, Figure supplement 3). Although the local diffusion coefficients are simulated to vary continuously within a certain range, just categorize the historical vectors of NP velocity into 3 clusters is enough to largely separate the regions of different binding affinity. As shown in Figure 5B, the 3 colors of blue, yellow and orange represent the NP velocity is high, medium and low, respectively. The segments where the NP undergoes directed motion is colored mainly blue. The regions where the NP binding occurs are colored mostly orange and yellow but of different portions, revealing their different binding affinity. Therefore, the combined SEES/MSD-ED and SEES/U-ED operations on the two observed variables (NP confinement and NP velocity) reveal both the locations and the affinity variations of the binding regions, providing a comprehensive description of the NP motion. We note that during the SEES/U-ED operation for velocity, if only 2 clusters are used, the different binding affinities in the confined regions will not be disclosed (Figure 5, red arrows); if 4 clusters are used, the redundancy on coloring the directed motions regions are clearly observed (Figure 5, purple arrow). Moreover, since the unsupervised k-means clustering is just capable of internal comparison, the obtained cluster centers could be randomly spaced and largely meaningless (Figure 5A, Row 5 in the left panel and Row 4 in the right). Only after the history-related inner differences are casted back to the trajectory, the NP dynamic states and their physical meanings can be identified.

**Figure 5.**
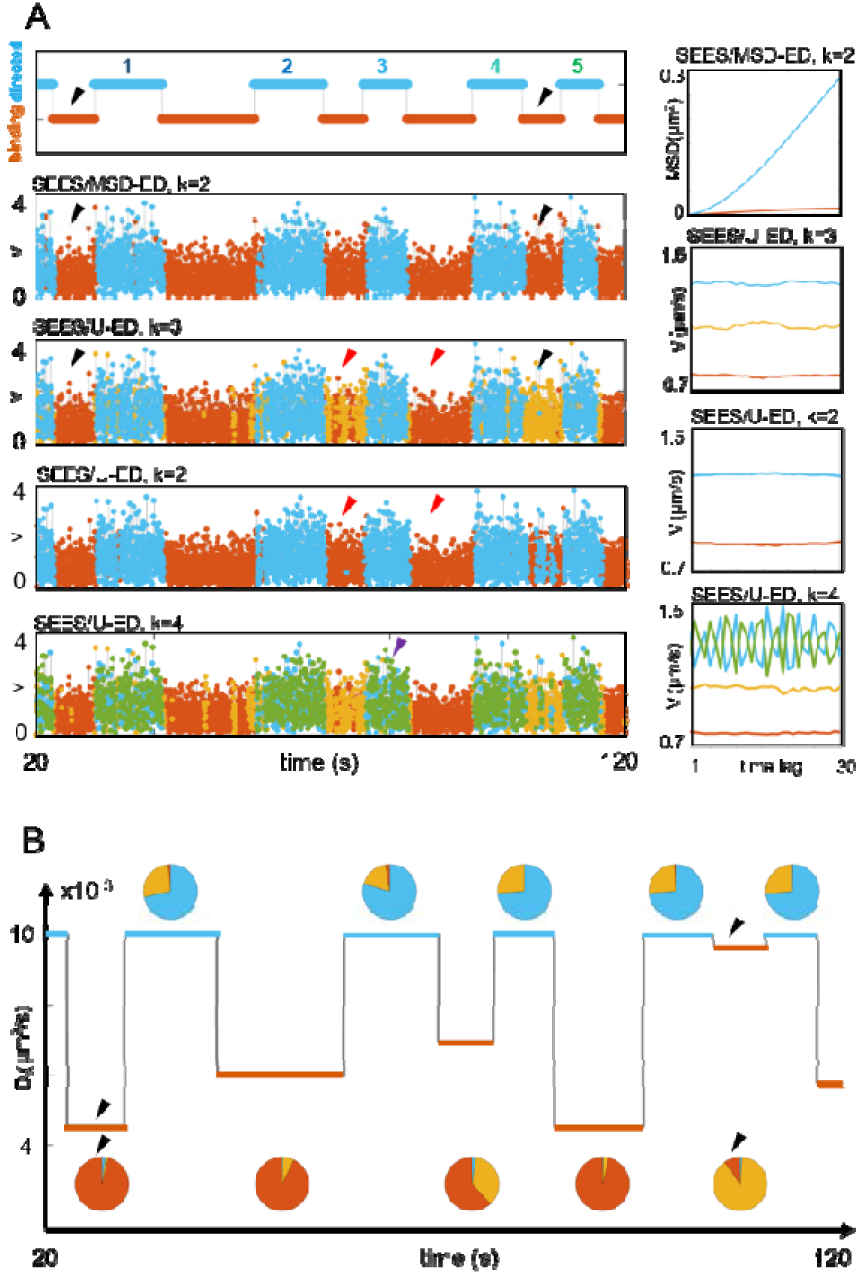
(A) Given the ground truth (row 1), the SEES/MSD-ED operator using 2 clusters differentiated the binding state and the directed motion state of the NP dynamics (row 2). The SEES/U-ED operator using 3 clusters further revealed the heterogeneity of the binding sites (row 3). Black arrows indicate two binding states with different local affinity. Row 4 and row 5 show the results given by the same operator with cluster number of 2 or 4, respectively. The cluster centers of these operations were displayed in the right panel. (B) The calculated local diffusion coefficients and the percentage distribution of the vector colors in each segments obtained from the SEES/MSD-ED operation. Black arrow indicates two binding sites with different local diffusion coefficient.

**Figure 6.**
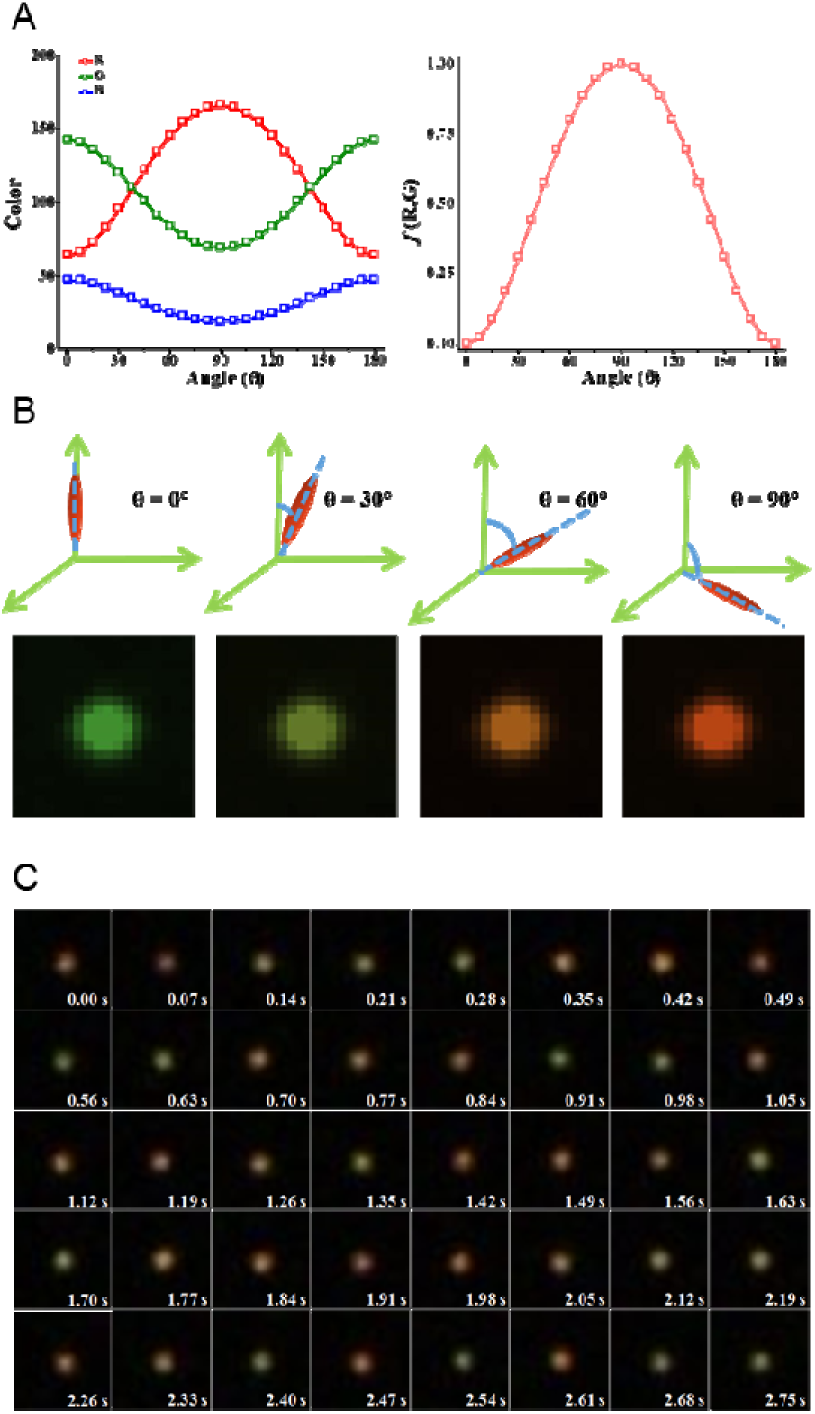
Out-of-plane angle of AuNR determination by color change. (A) Simulated darkfield image intensities of single AuNR obtained from the R, G and B channels of the color CMOS camera as a function of the out-of-plane polar angle θ and the polar angle dependence =sin2θ. (B) Typical AuNR polar angle orientations and the corresponding simulated color images. (C) Typical image sequence of a single AuNR rotated in 99.5% glycerol-water solution.

Altogether, though the physical pictures of the above 3 simulations differ considerably, the underlying NP dynamic states and state transitions were all successfully recovered by using the model-free SEES operator. A previously published dataset on cell migration dynamics (15) is also evaluated effectively (Appendix 1 Note S5). On the other hand, when applying the SEES operator to the simulated trajectory of a purely random walker with no motion state variation, no self-aggregated color segments is observed and different points on the trajectory are colored randomly regardless the length of historical experience vectors (Figure supplement 4). Therefore, the coloring profiles resulting from the SEES operator can only came from the real differences in the particle dynamics embedded in the particle trajectories. Since the SEES operation is capable of handling not only classical heterogeneous processes with just one temporal variable, but also more complicated processes with multiple time-dependent parameters and different physical meaning, this strategy may provide a new option for pre-processing of the multivariate big data.

## 4. Experiment: Single Nanoparticle Dynamics on Live Cell Membrane

To investigate the dynamic interactions between nanoparticles and cells, we monitored the entire transmembrane process of single gold nanorods (AuNRs) and resolved the corresponding diffusion dynamics with this model-free strategy. The SPT experiment was performed under a darkfield microscope equipped with a color CMOS camera. Because the diffusive and rotational behaviors are key parameters that can affect the nanoparticle fate in cell membrane (11, 32), we have developed a simple and effective method to simultaneously track translational and rotational motion of single AuNRs at the live cell surface based on the analysis of particle localization and out-of-plane angle-dependent scattering color (33).

Figure 7 displayed a complete trajectory of a RGD-modified AuNR entering the HeLa cell, which spanned nearly 9 minutes with over 8,000 data points. Intuitively, the spatiotemporal map could be divided into 5 local diffusional regions connected by 4 directional tracks (Figure 7A insert). To analyze the data, the conventional approach is to obtain the distributions of the AuNR angle, velocity and diffusion coefficient according to established physical models at the atomic-point or the sub-trajectory level with artificial segmentation or sliding window segmentation (Figure 7). According to the multi-parameter information as well as the predefined criteria based on their variations, certain dynamic patterns, temporal segments or critical points of the particle motion can be identified and used to describe the behaviors of the transmembrane process (Table1). However, the raw AuNR trajectories recorded in the heterogeneous cellular environments are highly complicated and are entangled with random noises, making it time-consuming to differentiate the dynamic states of the AuNR by studying the motion path of the particle visually even for trained technicians. As a result, personal experience based segmentation of the trajectory often leads to much subtle information unresolved or unobserved.

**Figure 7.**
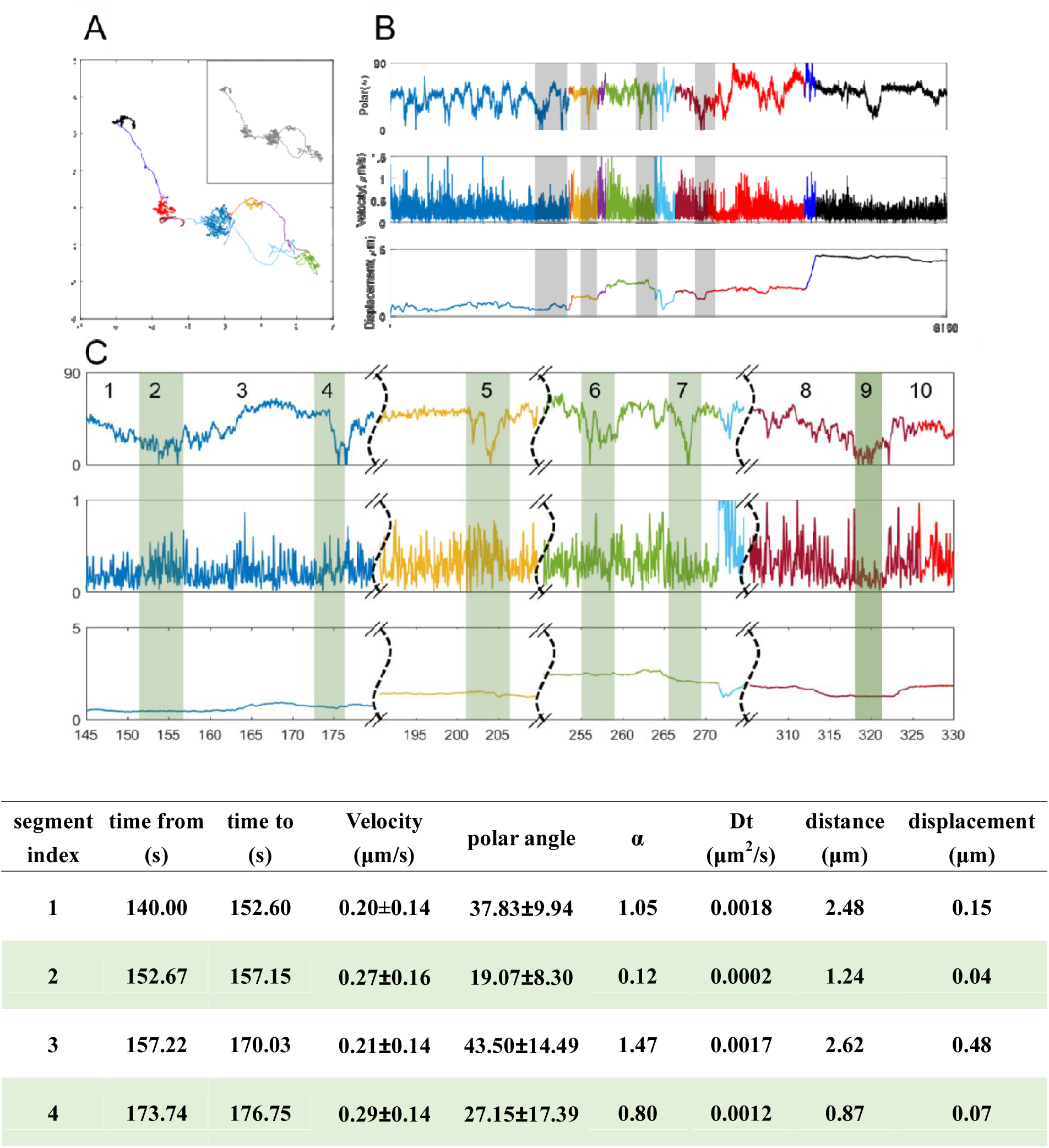
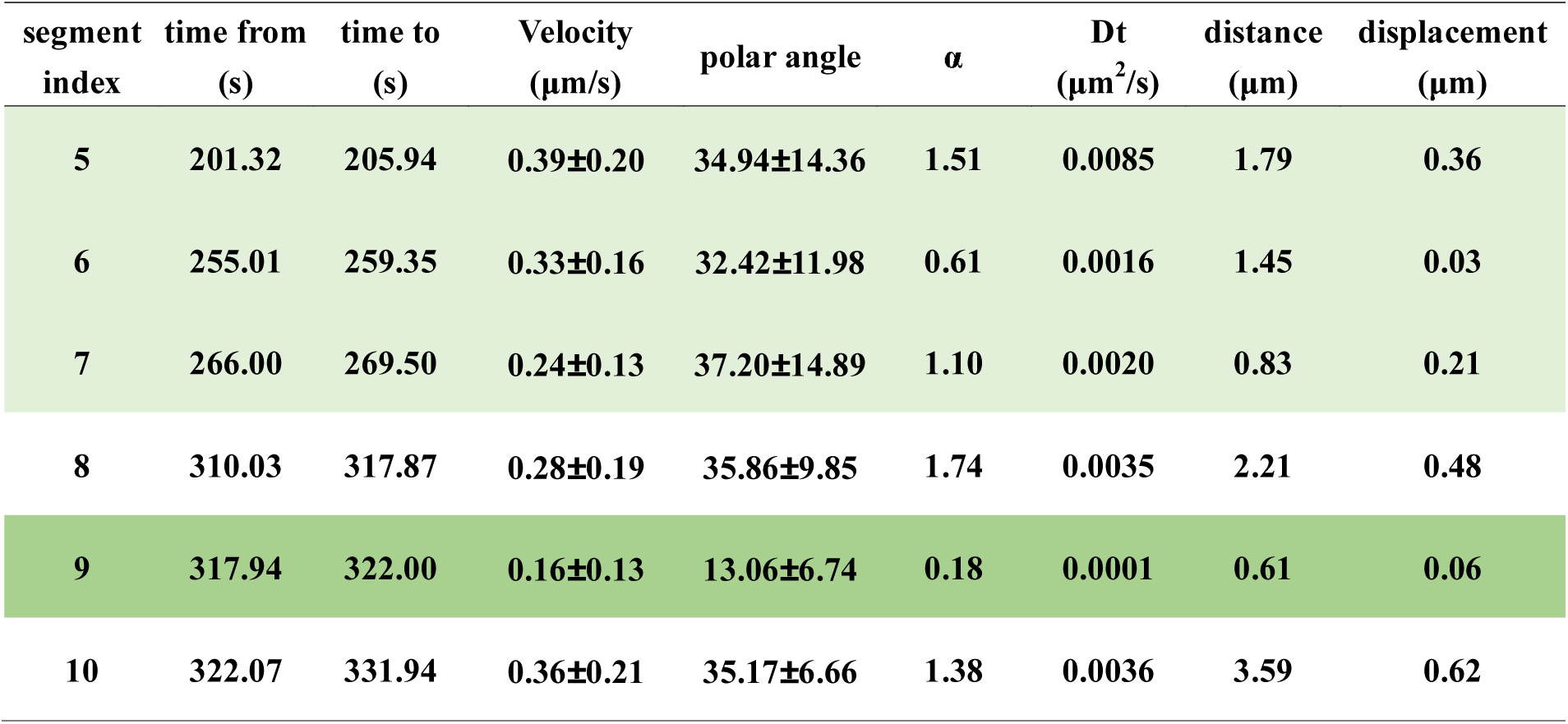
Traditional manual trajectory segmentations and cell-entry point determination. (A) RGD-coated AuNR interacted with HeLa cell membrane. Heterogeneous movement dynamics can be investigated by manual segmentation referring to the (B) polar angle, velocity and displacement time series profile. In this case, 6 diffusion states are linked by 4 directed states. (C) 10 regions were manually selected and their local physical properties were calculated as listed in table below. The cell entry site (region 9) was characterized with lowest mean velocity, lowest mean polar angle and confined movement.

Alternatively, the SEES method focuses on the variation trend associated with each spot rather than the atomic values, and are resistant to stochastic noises and local fluctuations. By processing the AuNR trajectory with the SEES operator, the segmentation of the NP trajectory and the classification of its motion states can be achieved directly through the resulting color-coded sequence, making it possible to analyze the diffusion statistics and reveal the transient heterogeneity of the NP-cell interaction dynamics. Figure 8 show the colored out-of-plane angle and velocity time-series after the SEES/U-ED operations. The experience-vectors were both categorized into 5 groups based on their magnitude from small (orange) to large (blue), respectively, resulting in the spatiotemporal trajectory being partitioned into tens of segments whose time duration varies from 1.33 s to 42.63 s. Figure 8C shows the accordingly colored x-y trajectory. The emergence of the sequence of color segments of various lengths produced a perceivable vivid overall picture on the temporality of the AuNR motion, which is not available from the raw or intuitively divided trajectory. When varying the length of the experience-vector (Movie S3) or trimming from 2 ends of the time-series (Movie S4), the overall coloring scheme from the SEES operation remains largely unchanged, as long as the vector length and the data size exceed some threshold. Therefore, the robustness and the reproducibility of the segmentation are guaranteed, and the SEES operation reveals true motion states of NP-cell interaction. For each segment, we can further calculate the feature parameters such as the average velocity, mean orientation angle, lateral and rotational diffusion coefficient, and anomalous factor of the particle during that period, respectively, according to established physical models. They provide transient NP dynamics and could serve as quantitative measures and statistical evaluation of the segments in the feature space. Such highly divided and uneven segmentation revealed the inner difference of the entire spatiotemporal process, which would not be achieved with artificial or sliding window subdivision.

**Figure 8.**
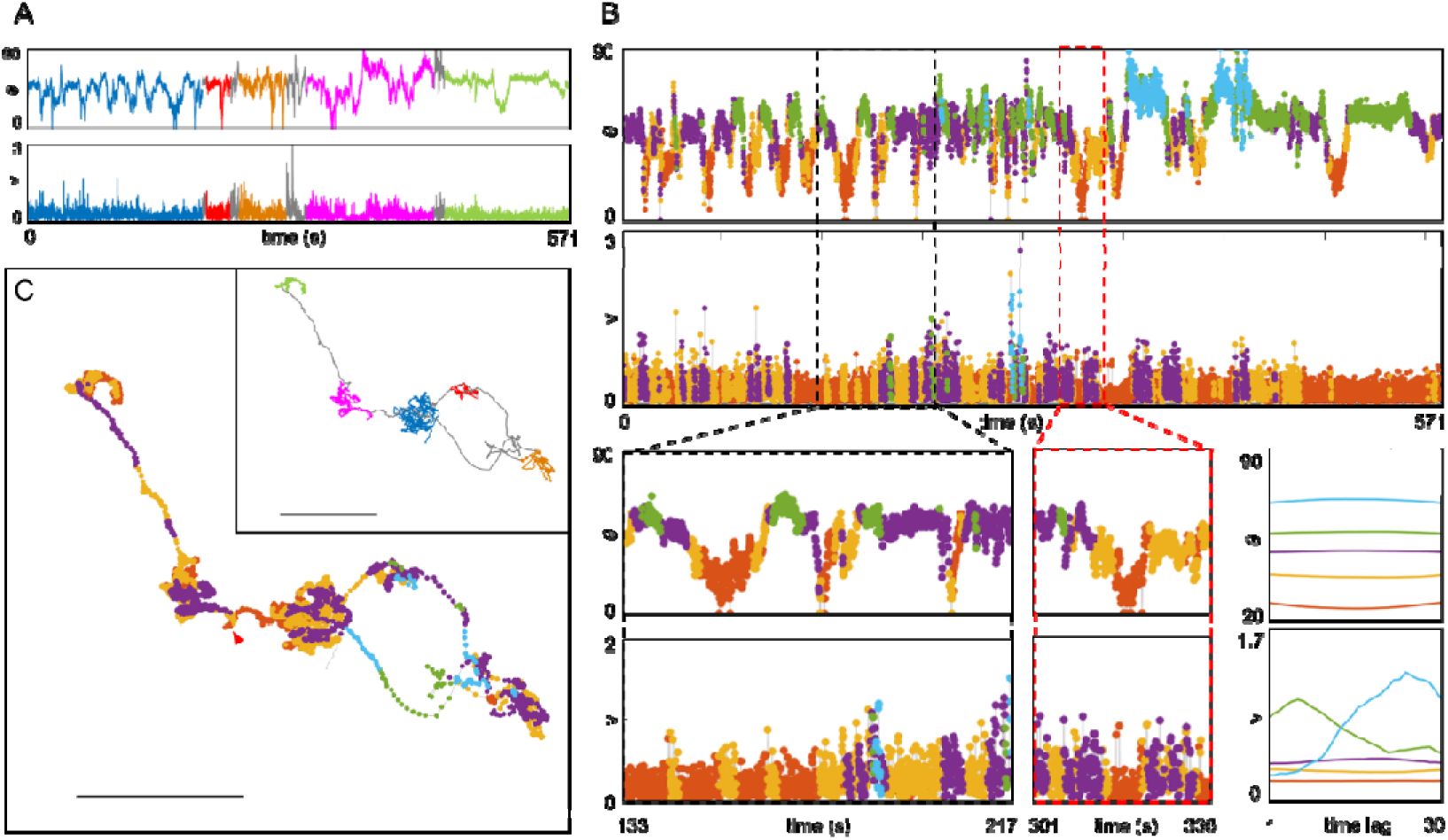
Identification of the single AuNR transmembrane site. (A) Time series of the out-of-plane angle (upper) and velocity (lower) of the entire dynamic process of a RGD-modified AuNR interacting with and crossing the membrane of a HeLa cell. The color segmentations are consistent with the manual segmentation of the spatiotemporal trajectory in the C inert. (B) Color segmentations of the same time series of the out-of-plane angle and velocity of the AuNR after SEES/U-ED operation. The enlargement shows that the transmembrane site (red dash box) can be easily differentiated from other similar sites (black dash box) by the synchronic tagging of orange color in both series. Also shown to the right of the enlargement are the cluster centers of the SEES/U-ED operation on the polar angle series (upper) and velocity series (lower). (C) The corresponding colored spatiotemporal x-y trajectory of the AuNR after SEES/U-ED operation on its velocity time series. The insert is the manual segmentation. The scale bar is 2 μm.

The SEES operation makes it easy to identify the spatiotemporal position of the transmembrane event. From the segmented out-of-plane angle and velocity time-series in Figure 8, it is clear that the translational and rotational motions of the AuNR near the cell surface are relatively independent in most situations. Nevertheless, the two types of motions became synchronous for a short duration at ∼ frame 4,500, displaying a same orange color region simultaneously. At this unique region, the AuNR out-of-plane angle and velocity were both small, indicating the nanorod at its highest confinement and lowest speed and being vertically inserted into the plasma membrane, which matched well with the reported situations of AuNR endocytosis (32). This cell-entry event was confirmed by the observation of directional transport of AuNRs after this region, which corresponds to molecular motor guided cargo transport after internalization by living cells. These findings suggested that the translational and rotational trajectory classifications under SEES operation make it easy for identification of the transmembrane sites of single AuNRs. By examining more single AuNR trajectories (Figure supplement 5 to 7), it turned out to be a correct measure in spotting the transmembrane sites and discriminate those particles that did not enter the cell, indicating the reliability of this data processing strategy. Furthermore, with the knowledge on the length of every tagged region, though only probabilistic, we could estimate the duration of single membrane-crossing events in an objective and reproducible manner. We determined the value to be ∼6.9 s for RGD coated AuNRs and ∼3.1 s cationic peptide coated AuNRs (Figure supplement 6 and 7). Therefore, the SEES segmentation provides a convenient method without modeling to determine the boundaries and transitions of single particle motion states in heterogeneous environments.

The SEES operation also allows in-depth investigation of the AuNR diffusional states and their transitions during the entire NP transmembrane process, which reflects the heterogeneous NP binding to the highly compartmentalized but fluidic plasma membrane as well as the crowding of the intracellular environment. To study the lateral confinement of the AuNR, we applied the SEES/MSD-ED operator to the AuNR x-y trajectory, where the sequences of experience-vectors were clustered into 5 groups according to the degree of local confinement (DLC) from high (orange) to low (blue) (Figure 9). As shown in Figure 9B, before the transmembrane event, the orange regions with the highest lateral DLC were relatively dispersed and their length was relatively short. After that, the orange regions spanned longer time and were more aggregated, indicating the AuNR mostly stayed in a crowded intracellular environment. A close examination on the chronological evolution of the colored segments (Figure 9C and Movie S5) and the historical vectors (Figure 9D and Movie S6, S7) discloses a more vivid picture. It is apparent that the transient diffusive behaviors of the RGD coated AuNR on the compartmentalized cell membrane could be divided into two types of hopping and lingering before the cell-entry, though the duration and displacement of either mode fluctuates all the time. The lingering length is typically several μm, where the particle was probably trying to enter the cell. The jumping length ranges from a few to tens of μm, providing more opportunities for the ligands on the AuNR surface to bind with membrane receptors of other locations once the transient ligand-receptor engagement was not sufficient for receptor-mediated nanoparticle endocytosis. Through frequent and drastic diffusive state transition, the particle could search a large area within a short time and find its appropriate cell-entry point quickly. We confirmed this motion pattern by analyzing the diffusion of multiple AuNRs. Such a hop-and-linger mode is consistent with that the plasma membrane is using two mechanisms jointly to adjust its local composition for membrane compartmentalization: the formation of barriers that passively hinder free motion and the active delivery of components to specific locations (34). Since it can achieve reproducible identification of changes in the diffusive behavior and reliable recognition of clustering and transient confinement within single particle trajectories, the SEES operator could make it more convenient to study membrane trafficking.

**Figure 9.**
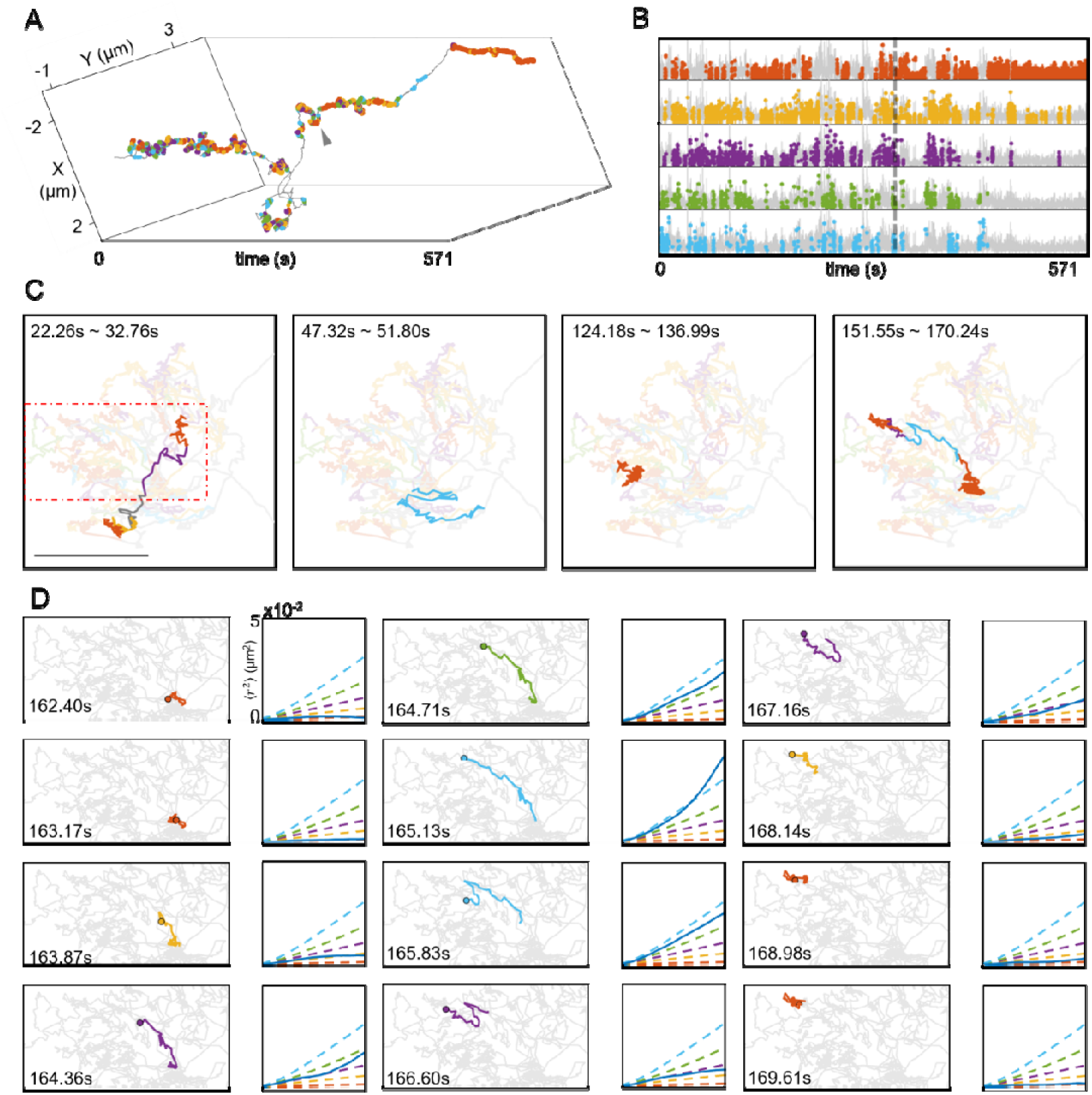
Single AuNR diffusional states and transitions revealed by SEES operator. (A) The colored 3D spatiotemporal trajectory of the RGD-modified AuNR in Figure 8 after SEES/MSD-ED operation on its velocity time series. (B) Temporal distribution of differently colored segments before and after the transmembrane site (dash line). The degree of confinement decreases from orange to blue. (C) Randomly selected 4 sets of segments show the hop-and-linger behavior of the AuNR as it searches for the cell-entry site in the first diffusive region (marked as blue in Figure 8A and 8C insert). (D) Sequential evolution of some of the experience vectors in the red dash box in C. For each pair of the graphs, the left one is the spatiotemporal trajectory of the experience vector carried by the time point; the right one is the MSD curve (blue solid line) as compared to the 5 cluster centers (dashed lines).

Besides analyzing one trajectory from different perspectives, the SEES operator could process multiple trajectories of different characteristics simultaneously. As an example, we compared the cellular uptake of single AuNRs coated with RGD and cationic peptide, respectively. The later takes an electrostatic adsorption mediated internalization mechanism that is different from the receptor-mediated endocytosis pathway. For 2 set of trajectories of each case, we used SEES/MSD-ED, SEES/U-ED and SEES/AC-ED operator to analyze the local spatial confinement, motion speed and rotation rate of the 4 particles, respectively. By counting the number of different color points on each colored trajectory (Figure 10A), we obtained 12 state frequency histograms representing the unique dynamic profiles of the single AuNR interacting with HeLa cell membrane (Figure 10B). While the cationic peptide-coated AuNRs covered a smaller membrane area, they have less lateral confinement, higher translation speed and faster rotation rate than the RGD-coated AuNRs, implying a smaller DLC dominated by weaker interaction between the cationic peptide and the cell membrane. RGD-coated AuNRs stayed longer durations at high DLC state and took more time to enter the cell. In contrast, cationic peptide-coated AuNRs displayed frequent transitions between high and low DLC states and spent less time on penetrating the cell membrane. These observations could be attributed to that the RGD-coated AuNRs suffer from gradually increased confinement due to multiple receptor-ligand binding before endocytosis, while the interaction between the cationic peptide-coated AuNRs and the negatively charged plasma membrane is governed by electrostatic interaction. The abundant information extracted from the multivariate trajectories could be beneficial to in-depth study of the diffusion dynamics of various nano-cargoes at the single particle level.

**Figure 10.**
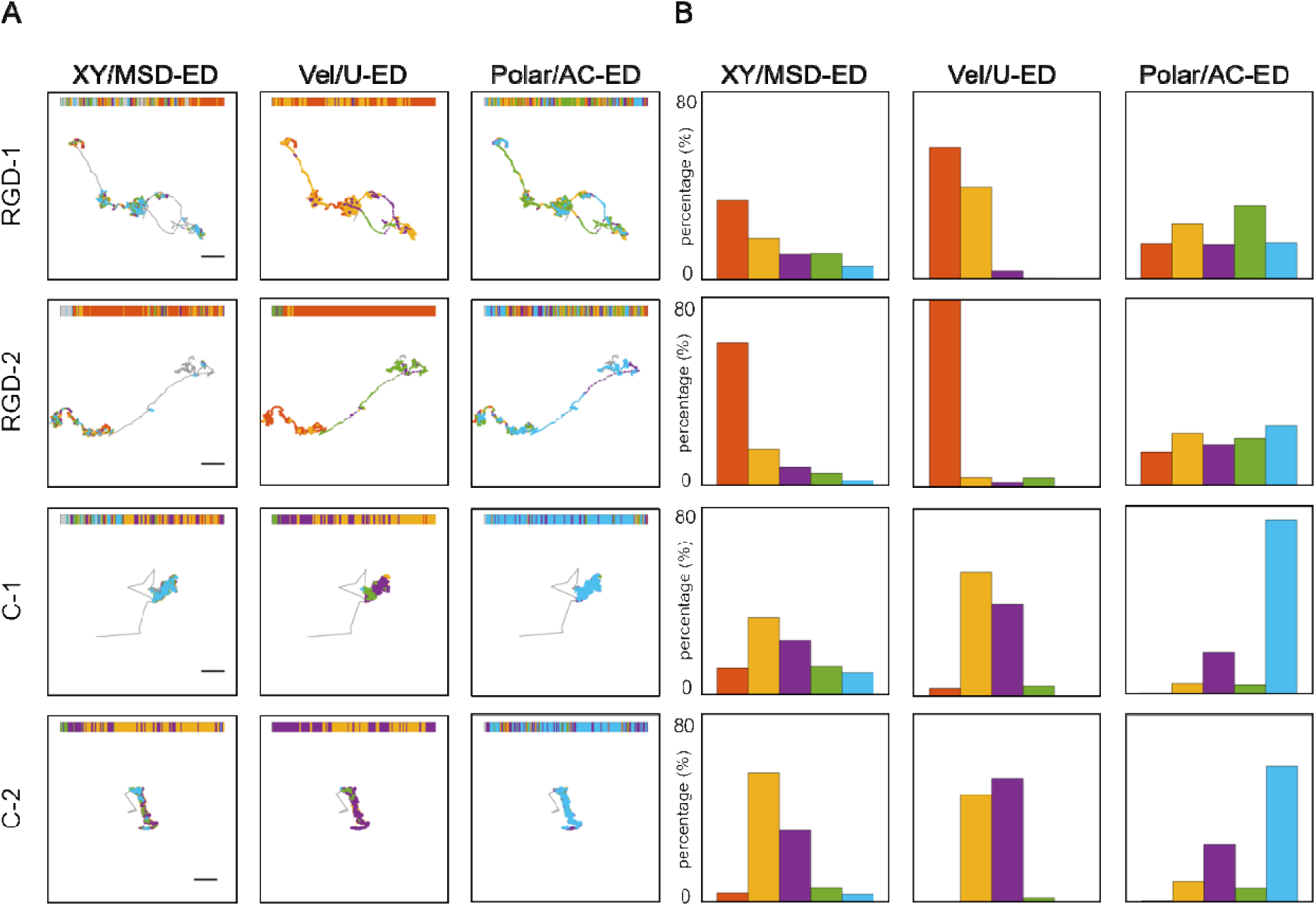
Differential dynamics of RGD and cationic peptide modified NRs. SEES/MSD-ED, SEES/U-ED, SEES/AC-ED operators were applied to analyze the spatial positioning, velocity and polar angle of 2 RGD-modified AuNR (RGD-1 and RGD-2) and 2 cationic-peptide-modified AuNR (C-1 and C-2) transmembrane dynamics, respectively. (A) Each box shows the colored segmentation of the corresponding time series (color bar code insert) and its mapping onto the spatiotemporal trajectory. The scale bar is 1 μm. The length of all trajectories used here was 8000 frame (560s). (B) The group percentage profiles revealed by the corresponding operator.

## 5. Discussion

We have demonstrated through both simulation and experimental data analysis that the SEES operator is able to uncover the hidden motion mechanism or the inner differences embedded within the observed dynamics. Adopting the AuNR-cell membrane interaction for a case-study, we show that a combination of particle velocity, rotation angle and spatial position data can be readily analyzed using the same format without predefining a set of models and parameters for each differently recorded time series. This multivariate SPT analysis provides a comprehensive visual interpretation of the particle dynamics, and the resulting color labeling and segmentation of the trajectories make it convenient to identify the boundaries, transitions and critical points of the motion states of various functionalized NPs.

More generally, the trajectory of an individual particle can be considered as a time-course evolution in state space and thus comprise not only widely studied spatial coordinates but also other variates that describe the dynamics of the particle such as scatter light intensity and color under the microscope. As the imaging equipment and tracking algorithm developed rapidly, more dimensions of the variates as well as larger data set are obtained from a single experiment. Although the pre-defined physical model is essential to interpret data with limited size, this strategy may face its difficulty of “model explosion” in dealing with multivariate trajectory in state space, that is, the number of models and parameters needed to fully describe a certain particle dynamic increase extraordinarily versus the number of observed physical parameter. The SEES operation could be a pre-processing step to simply let the inner structure of the time series emerge by themselves without prior assumptions of the system, and therefore provide with both an intuitive picture and quantitative characterization of the multivariate trajectory by the same algorithm. The emergence of an intuitive color picture of the whole system with minimal subjective bias, combined with prior knowledge on the particular system, would facilitate the formation of new hypothesis, interpretation and models. The abundant information obtained from this presuppositionless method without averaging, fitting or oversimplification could be further used to select the critical physical parameters for modeling and inspire new knowledge finding, or downstream biological/physical evaluation. Therefore, this method could help studies on heterogeneous individual dynamics, especially those lacking prior knowledge or multiple variates being observed. The uniqueness of the SEES operation lies in the “negation” of the mechanical viewpoint on the acquired spatiotemporal data. Rather than viewing the data as mere assembly of isolated points, frame them with external *a priori* models, and try to establish an “objective description” of the complex system, we assign a short history to every data point, differ them based on the variation of their history, and tag them according to their classified history. Just like labeling protein or DNA biopolymers with fluorophores, coloring single particle trajectory with its own history exhibits a comprehensible representation of the original spatial or spatiotemporal structure. On the other hand, unlike static fluorescent labeling, dynamic historical labeling of a single particle trajectory is not exclusive. How to construct the history of the data point and how to compare the history determines how to color the trajectory. Nevertheless, as long as the SEES operation is reproducible, the resulting coloring profile is the truth of the spatiotemporal difference of the particle behavior, and the diversity of color labeling profiles reflects the complexity of particle dynamics and its surrounding environment. After all, no data would be meaningful if not being interpreted appropriately. Wherever relying on external models or internal historical comparison, some sort of subjective-objective unification is necessary during any data processing.

Besides analyzing the single particle trajectories, this data-phenomenology strategy provides a general framework for spatiotemporal series analysis of complex systems aiming to reveal their interconnectivity and dynamic evolution. For decades, mechanical-model-based data-analysis and computer-simulation have played significant roles in advancing our understandings on the fundamentals and the prediction capability of system behavior in numerous research fields. On the other hand, the arbitrary division between the essential laws and the changing phenomena often restricts researchers from going deep into the water of complexity. Our approach, with no predefined external models and laws, provides a novel rationale that allows the underline mechanisms of complex systems to emerge on their own out of the otherwise huge amount of clueless data. The SEES operation meets the recent trend “to let data speak for themselves” in data science and could potentially inspire the finding of new knowledge of many systems. We expect that the unification of the phenomenology approach and data science will fundamentally promote the studies of the complexity issues.

## Appendix 1: Supplement Notes and Figures

### Note S1. The Theoretical Foundation of the SEES method

In the main text, an analogy to the listening to the music is used to introduce the SEES method. However, its theoretical foundation roots in the phenomenology of the notion of time, i.e. how we experience time or how to account for the way things appear to us as temporal. Here, the problem we face is to describe a single particle trajectory as a temporal object. Obviously, we would not analyze any trajectory acquired using the microscope but only those appear to us carrying meaningful information, therefore, we must have intentionally or subjectively assumed that there is some useful unity as well as details embedded in the trajectory before performing the analysis. In that, analyzing a single particle trajectory through some computer algorithm is just like listening to a music through the structure of our consciousness, and the difficulty is well-illustrated in Kelly’s article (https://www.iep.utm.edu/phe-time/):

> “To highlight the difficulty and importance of explaining the structures of consciousness that make possible the experience of time, Husserl, like his contemporaries Henri Bergson and William James, favored the example of listening to a melody. For a melody to be a melody, it must have distinguishable though inseparable moments. And for consciousness to apprehend a melody, its structure must have features capable of respecting these features of temporal objects. Certainly, we can “time” the moments of a temporal object, a melody, with discrete seconds. But this scientific and psychological account of time, which, following Newton, considers time as an empty container of discrete, atomistic nows, is not adequate to the task of explaining how consciousness experiences a temporal object. In this case of Newtonian time, each tone spreads its content out in a corresponding now but each now and thus each tone remains separated from every other. Newtonian time can explain the separation of moments in time but not the continuity of these moments. Since temporal objects, like a melody or a sentence, are characterized by and experienced as a unity across a succession, an account of the perception of a temporal object must explain how we synthesize a flowing object in such a way that we (i) preserve the position of each tone without (ii) eliminating the unity of the melody or (iii) relating each tone by collapsing the difference in the order between the tones. Bergson, James and Husserl realized that if our consciousness were structured in such a way that each moment occurred in strict separation from every other (like planks of a picket fence), then we never could apprehend or perceive the unity of our experiences or enduring objects in time otherwise than as a convoluted patchwork…”

It is not difficult to see that in the above citation, if we replace the word “melody” with “trajectory” and “tone” with “data point”, all the reasoning still holds. Because previous SPT analysis methods are all based on the Newtonian time, in order for us to avoid subjective bias and experience the single particle trajectory as a unity across a succession, the key issue is to find a solution that would not treat the trajectory as a sequence of discrete, atomistic nows in an empty time container. While different philosophers have established various theories, herein we favor the dialectic of time or the negation of negation of now (this moment) articulated by Hegel. In the first chapter (Sensuous-Certainty; or the “This” and Meaning Something) of his book Phenomenology of Spirit (ξ106−ξ107), Hegel said:

> “… the Now is just this Now as it no longer is. The Now is, as it has been pointed out to us, what has been. This is its truth… It is nonetheless true that it has been. However, what has been is in fact no essence; it is not, and the issue at stake had to do with what is… In this pointing out, we therefore see only a movement and the following course of the movement. (1) I point out the Now, and it is asserted to be the true. However, I point to it as something that has been and thus sublate the first truth, and (2) I assert the Now as the second truth, that it *has been*, that it is sublated. (3) However, what has been is not; I sublate that second truth, that it *has been*, or, its having-been-sublated, and, in doing that, I negate the negation of the Now and so turn back to the first assertion, namely, that *Now* is. The Now and the pointing out of the Now are therefore composed in such a way that neither the Now nor the pointing out of the now are what is immediately simple. Rather, they are a movement which has various moments in it; *this* is posited, but instead it is *an other* which is posited, or the This is sublated, and this otherness, or the sublation of the first, is itself *again sublated* and in that way returns back to the first. However, as reflected into itself, this “first” is not wholly and precisely the same as what it was initially, namely, an *immediate*. Rather, it is just something reflected into itself, or a simple which remains in otherness what it is, namely, a Now which is absolutely many Nows, and this is the genuine Now, or the now as the simple daytime that has many Nows within it (that is, hours). Such a Now, an hour, is equally many minutes, and it is this Now which is equally many Nows, etc. – *Pointing out* is thus itself the movement that declares what the Now in truth is, namely, a result, or a plurality of Nows taken together; and pointing out is the experience that the Now is a *universal*.”

In brief, without *a priori* definitions, the Now can only be defined by what are not the Now, that is, to characterize the Now with its adjacent historical points. From the “succession of experiences” to the “experience of succession”, the whole procedure performed by the 3-step SEES operator on the single particle trajectory is essentially the “negation of the negation” of the momentary movements of the particle. Unable to rely on predefined models but still taking the entire trajectory as a unity, an individual moment can only find its relative “truth” via making active connections to all others. Making an argument similar to Hegel’s dialectic. Firstly, to construct the historical experience-vector is the “negation” of atomicity of the individual “now” moment, and disburses its zero dimensional information into a high dimensional space. The appearance of every moment in continuous tens of vectors and the heavy overlap between adjacent vectors ensure the plurality of the “negation” action and make them all connected. Secondly, the unsupervised clustering, carrying a particular predisposition from the observer, drives each individual vector to compare with all others universally and find out their relative position by using other vectors as “mirrors”. Finally, by coloring every momentary point on the NP trajectory according to the category of its leading vector, the first “negation” is “negated” again, and the added information from vectorization and universal comparison is “reflected” back into the original individual moment itself. The high dimensional space is reduced back to the low dimensional space, and the global clustering difference is projected onto the localized spatiotemporal difference. Such a dialectic circle, from the momentary individuality to the universality of the whole trajectory and then back to the individuality, is achieved through the particularity of all the individuals’ own history rather than through externally defined mechanical models. Consequently, a continuous “change of change” of the single particle movement both globally and locally is exhibited in the form of a colored trajectory, which is just like the details as well as the unity presented aurally by a music. To our knowledge, it is the first time that the “negation of the negation” dialectic principle, supported by the machine learning technique, finds a practical implementation in fundamental scientific research. Alternatively, since extensive vectorization, universal comparison, and certain reflection are the common aspects of almost all machine learning techniques, the “negation of the negation” dialectics probably underlies the effectiveness of the methodological foundation of the Data-Science.

### Note S2. The Description of the SEES operator

The algorithm of the SEES operator implemented by MATLAB script mainly consists of three parts, the construction of the historical experience vector, the unsupervised clustering and results viewer based on the color code mapping corresponding to the three steps described in the manuscript. For time series observations of a particle dynamics:

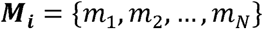

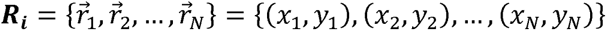

The ***M_i_*** is the time series observation such as velocity or rotation angle while the ***R_i_*** represents the time series of the particle locations. Time intervals between consecutive observations is Δt. The historical experience-vector can be constructed by 3 means provided by the algorithm:

1. Univariate method

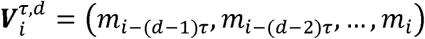

2. MSD method

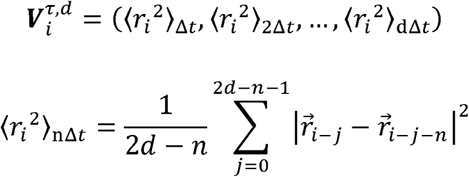

3. Autocorrelation method (AC)

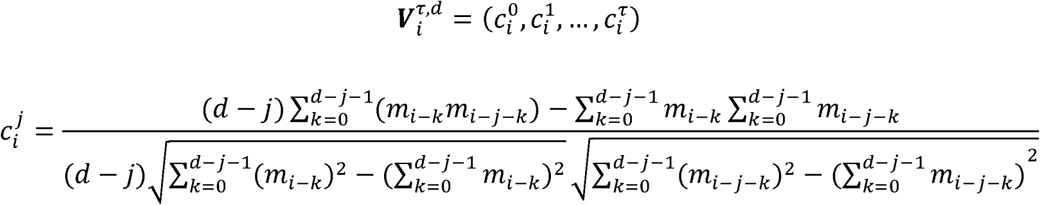

Here, 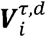 represents a historical experience-vector constructed in the first step and anchored at a specific time point *i*. For each construction method, an initial offset is required as the beginning of the series, where the historical experience detected is shorter than the length of the vector and hence invalid. The offset for the univariate, MSD and AC methods are 1 + (d − 1)τ, 2d and d respectively. The entire sequence of the constructed experience-vectors can be represented as a matrix ***W*** and is ready for the following unsupervised clustering.

The univariate and AC method are suitable to deal with single time series variable ***M*** while MSD method is meant to handle spatial movement series which has the two ***X*** and ***Y*** coordinates. Univariate method is helpful to extract historical experience variation values in the time series, while the AC method mainly focuses on the fluctuation properties of the series. The MSD method is suitable to investigate the historical confinement experience as its physical definition indicates.

In the next step, the experience-vectors are compared globally to let the internal differences emerge. Here, to prove the concept, we adopted the k-means clustering, which is one of the most widely used unsupervised clustering algorithms, as our classifier. Similarity measurement can be performed with various metrics to focus on different features of the vector. The metrics we adopted here include Euclidean (ED), Cross-Correlation (CC), and Minkowski Distance (p=5, MD).

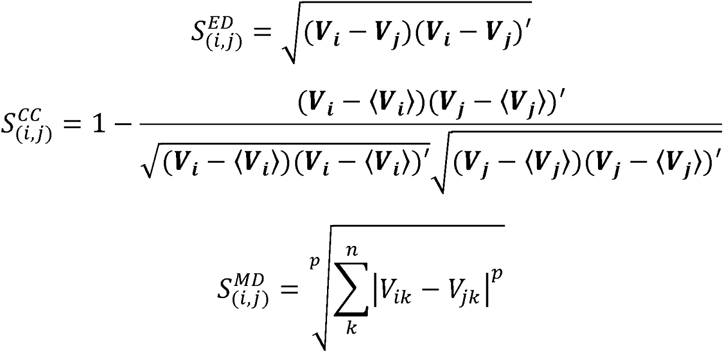

The metric selection helps us to focus on different aspects of the experience vector. CC metric is sensitive to the variation trend of the sequence of elements in the vector, while the ED and MD metrics mainly focus on the magnitude of the vector elements. As the MD is a generalized form of the ED, their clustering results are often comparable to each other. In MD metric we set p=5 as an odd number to make this metric more robust to fluctuations. To meet more specific needs, more advanced metric such as DTW^1,2^ can also be adopted in the clustering step. Considering both the experience vector construction and the metric selection, theoretically we can provide 9 different combinations in the algorithm. With the univariate method (U), the U-ED/U-MD combination is capable of differentiating the magnitude of experience vector of a specific measurement, such as the velocity or angle variation of the AuNRs. Meanwhile the U-CC combination is suitable to differentiate the historical tendency like what we did in the double-sinusoidal model in Figure 2. AC curve usually works with ED metric to differentiate the historical fluctuation behavior such as the rotation dynamics of the AuNRs as it always exhibits similar decade tendency. MSD-ED/MSD-MD consider the magnitude of historical mean displacement versus the increasing time interval while MSD-CC only consider the tendency of the curve growing.

In this work, we use the basic and well-studied k-means method to perform the clustering. Again, more advanced clustering methods or cluster number estimation method can be applied in this step. Here, we optimize the performance of the classical k-means by two ways. First, multiple runs were used to reduce the interference due to the random initiation, optimized result was selected by the minimal sum of sample-cluster-center distance. Meanwhile, rare cluster whose population is lower than a threshold (usually 10%∼20%) was marked as undefined group to ignore the effect of outliers.

After classification of the historical experience-vectors, the original spatiotemporal trajectory or the time-series of the AuNR velocity or angle were colored according to the clustering results. Besides providing an overall color segmentation pattern, an algorithm is designed to identify the exact frame-wise positions of each same color segment. To eliminate too short noise segments, we set up a 20-frame-length (1.4 s) threshold. A segment shorter than 20 frames would be combined into the following one until its length reaches 20-frames. Several visualization methods were implemented in MATLAB as well to enable the researchers to navigate the spatiotemporal series step by step and get an intuitive picture of the collective dynamics.

The main function of the MATLAB codes is pNPa which returns a NPMotionTest instance containing all the data-processing results. The visualization was performed by PNPGUI class which accepts a NPMotionTest instance and then generate a user-friendly GUI interface. The user can easily navigate around along the whole trajectories to catch the dynamic details as well as the spatiotemporal patterns as a whole. The calling formats of these functions are:

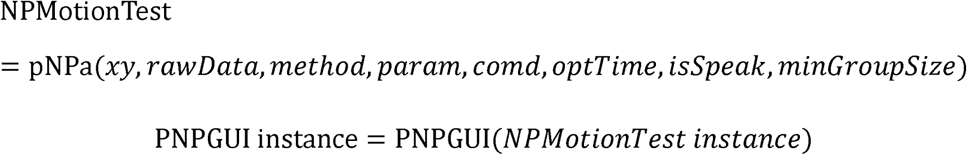

More description of the function options can be found in the table below.

Parameter descriptions of the functions.

Parameter descriptions of the functions.
| Input parameter | Explanations |
| --- | --- |
| *xy* | particle trajectory formatted as **N×2 matrix**, each row is a XY coordinate observed at specific time |
| *rawData* | processing raw data, the valid format depends on the *method* (Table 2) |
| *method* | the method for historical vector extraction, a **String** variable |
| *param* | a 1×4 vector parameter for historical vector extraction, see Table 3. |
| *comd* | cluster metric, a **char** variable. **E** for Euclidean Distance, **C** for Corss-Correlation and **M** for Minkowski Distance. |
| *optTime* | cluster repeating time. Each time the algorithm converges in a local optimization. Repeat with randomly initial state helps to obtain global optimization, a **scalar number**. |
| *isSpeak* | is plot the result at the end of processing, a **logical** variable |
| *minGroupSize* | constrain the minimum number of group members in cluster to avoid the outliers effect. a **fraction** number. |

### Note S3. Model simulation

1. The rotation/revolution model. The rotation/revolution model was simulated by combining two sinusoidal functions with different frequencies.

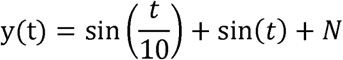

Here, the ***N*** represents a standard normal distributed random noise.

2. Random walk on heterogeneous 1D cell membrane. A virtual 1D cell membrane was set ranging from -∞ to +_∞_ and served as the default region. Then, several specific local regions were added with heterogeneous properties. The local properties of a specific region of the simulated cell membrane consist of the local diffusion coefficient ***d_t_*** and direction preference ***p***. The direction preference provides a bias of the direction selection in random walk. *p* ∈ [−1,1} where −1 means absolute negative movement and +1 means absolute positive movement. The step length was generated as following:

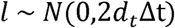

where Δt is the time interval of the simulation. The particle was placed at a specific coordinate and then allowed to diffuse based on its local regional properties. The expectation of the particle velocity is given by:

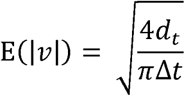

3. State switching on 2D cell membrane. The 2D cell membrane was generated with an enclosed boundary and a specific diffusion coefficient *d_t,0_.* Then, a number of non-overlapped binding sites were placed randomly on the cell membrane. For each binding site *i*, four parameters were used to specify its binding behaviors: confined radius *r_i_*, local diffusion coefficient *d_t,i_*, affinity coefficient *a_i_*and escape probability *p_i_*. The particle initiated on the membrane with Brownian motion. After entering a certain binding site, the particle can alter its state to the binding state according to the affinity coefficient of the site. In binding state, the particle movement in each step was divided into 100 sub-steps and make sure that the particle is confined inside the binding radius. After each confined step, the particle can take a chance to release itself and transform to directed motion according to the escape probability. The motion step length was generated according to the density function^3^ F*_l_*:

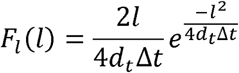

The directed motion was simulated by direction restriction:

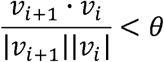

### Note S4. Meaningfulness evaluation on 3 simulated trajectories

As the subsequence time series (STS) clustering was proved to be meaningfulness in Keogh’s paper^4^, one may raise the question of whether the preprocessing of SEES operator is meaningful, especially when the Univariate vector construction seems to be similar to the operation of STS clustering. To answer this question, firstly, we note that rather than focusing on the physical meaning of the cluster centers as in Keogh’s paper and the conventional time series clustering methods, our SEES operator, as a preprocessing method, focuses on revealing the temporal coloring pattern along the trajectory. Secondly, we designed a computation experiment to show how the SEES operator successfully uncover the underlying pattern of the simulated time series (shown below). The 3 steps of the SEES operator (experience vector construction, unsupervised clustering and coloring the trajectory with group tag) together as a whole make up the SEES method, and our method cannot be reduced to any of the individual step.

We have designed a simple but intuitive computational experiment to evaluate our method. A total random series ***x1*** was generated following the standard normal distribution. Next, we sorted the ***x1*** series with increasing magnitude and generated series ***x2***. Finally, we cut the ***x2*** series into 3 pieces and shuffled the piece order and the element order in each piece and generated series ***x3***. From ***x1*** to ***x3***, we simulated 3 series with identical elements but different underlying mechanism (Figure A below). Following the experiment designed in Keogh’s paper, we applied the SEES operator to each of the series with univariate experience vector construction and randomly restarted 3 times for each. The typical results are shown in Figure B below. As expected, the random series ***x1*** shows random pattern and its related clusters centers seems meaningless. However, the SEES operator captured organized temporal pattern of sorted series and shuffled series with different magnitude of clustering centers.

We also calculated the ‘meaningfulness’ of the computation following the formula in Keogh’s paper.

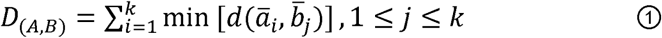

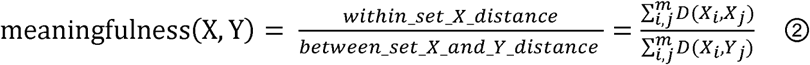

Note that the meaningless claimed in Keogh’s paper means that the cluster centers from different data set have similar (or identical) pattern. *D*_(*A,B*)_ is a distance measurement of two cluster centers, A and B, which consists of k cluster centers 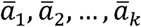 and 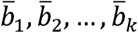. *d*_(*a,b*)_, respectively, are a distance metrics of vector a and b, such as Euclidean Distance, etc.. X and Y are two different time series. _i_ is the cluster centers given by i*th* initial start of k-means clustering, total number of initial start is m. We found that although the cluster centers of random series and sorted series were largely different, but the meaningful factor appeared high in our calculation, which means that the cluster centers between the two sets were similar to each other (green bar in Figure C below, left panel). We found that the cluster distance formula ① presented in the Keogh’s paper didn’t forbid the case where all centers in one dataset match to the same center in the other set, and thus cannot fully reflect the differences between two groups of cluster centers. We then changed the cluster distance formula from simply adding up minimal distance between cluster centers to a dynamic programming algorithm (Figure C below, right panel), which guarantee the cluster centers between two sets were matched one-by-one. The modified formula gives more reasonable results that the clusters results among the 3 series are different from each other. Note that this result can be different for each time of restart because the multiple restart optimization algorithm of k-means is forbidden according to Keogh’s paper. In summary, the key point of our method is the emergence of a temporal color pattern instead of the cluster centers, and we proved that the SEES operation can give correct patterns in typical tasks.

**Note Figure 1.**
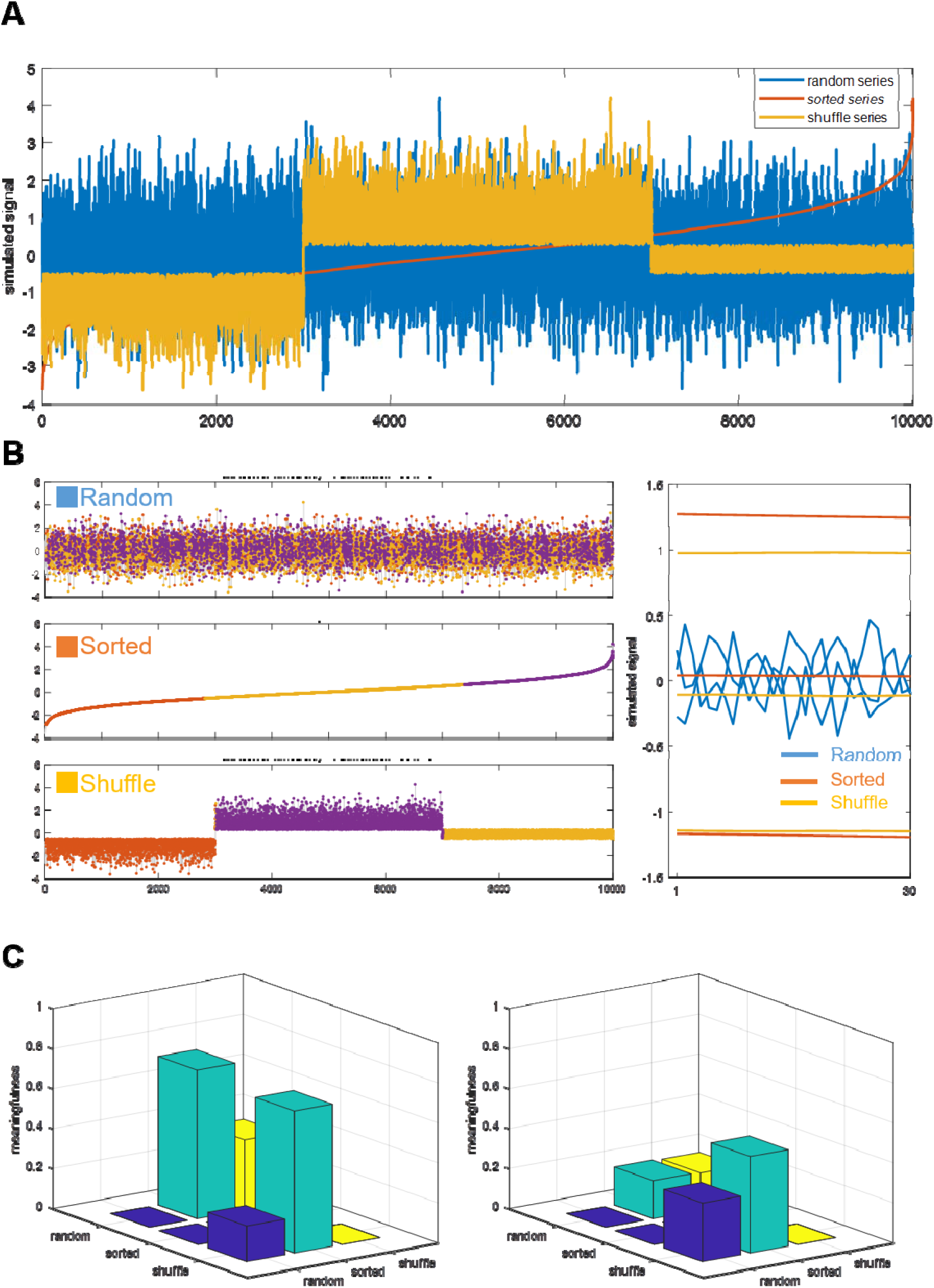
(A) Three time series were simulated with the same set of numbers. (B) SEES/U-ED was applied to each of the series with 3 clusters. (C) Meaningfulness calculated by Keogh’s formula (left panel) and dynamic programming (right panel).

### Note S5. Revealing the underlying pattern of cell migration trajectories from a previously published paper

To evaluate the reliability and wide applicability of our method, we applied the SEES operator to the simulation data of a published paper^5^, in which a persistence-active model was proposed to analyze the cell migration trajectory. The model is based on the auto-regression statistics and has two terms to describe that cell migration dynamics, the activity term and persistence term.

Leaving the underlying model and mechanism unknown, our method provided reasonable coloring pattern comparing to the ground truth. As shown below, for the 3-steps staircase regime-switching case, the univariate experience vector of velocity (U/Vel) was constructed and grouped in 3 classes. The projected color roughly reveals the 3 segments structure (Figure A below). Besides, the autocorrelation experience vector of X displacement (AC/X) also helps to uncover the same structure (Figure B below). Interestingly, we found that the velocity series was more sensitive to the active term while the particle displacement was more sensitive to the persistence term of the model, which suggested that multivariate time series analysis enabled by our pre-suppositionless approach was capable of revealing the system dynamics from different perspective. Note that the U/Vel setup gives a noisier result for the intrinsic randomness of the model (normal distribution term).

The second case is a continue-changing state with unsynchronized persistence-active dynamics. As a result, the U/Vel and AC/X setup of the method also revealed different underlying dynamics. U/Vel that is sensitive to active term shows a growing pattern which is consistence to the ground truth (Figure C below). One the other hand, the AC/X that is sensitive to the persistence term exhibits the symmetric pattern as the persistence grows firstly and goes down at last in the ground truth (Figure D below). The clustering centers of U/Vel and AC/X also correctly shows the increasing (from red to purple) and the anti-self-correlated (red and yellow) /self-correlated (purple) pattern, respectively.

In summary, our method successfully reveals the underlying dynamics of the evaluation data in previous published paper while no prior-knowledge of the system or model is involved in the data processing. As the underlying mechanism of cell migration is largely different from that of the Nanoparticle diffusion, the results indicates that our method is reliable and potentially applicable for wide research field based on single particle tracking.

**Note Figure 2.**
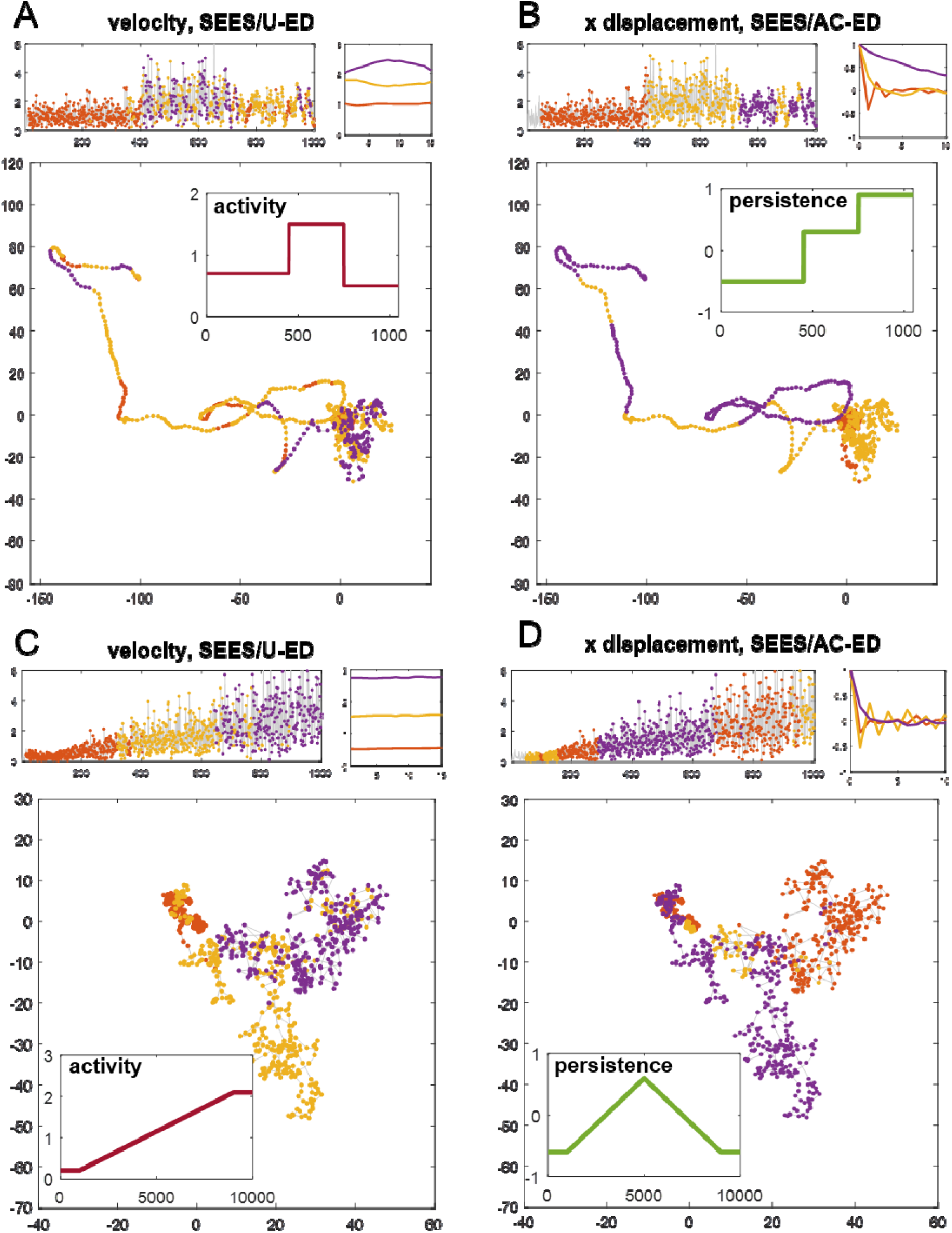
SEES/U-ED and SEES/AC-ED operators were applied to velocity (A) and X displacement (B) time series in three steps staircase mechanism, respectively. They were also applied to the velocity (C) and X displacement (D) time series in the continue-changing mechanism. Note that the similar results can be obtained with Y coordinate compared with those given by the X corrdinate.

### Supplementary Figures

**Figure supplement 1.**
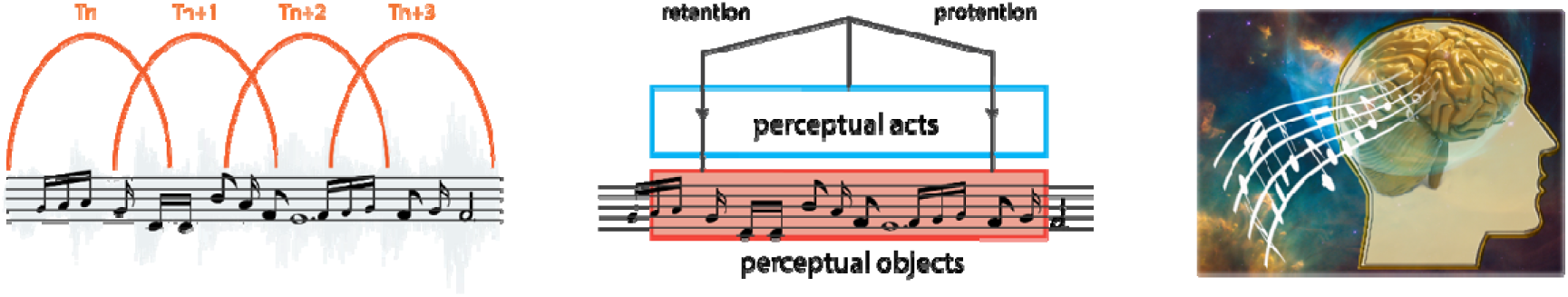
The phenomenology of time-consciousness, using enjoying music as an example. (left) A melody can be represented as a collection of isolated notes. At each time point, however, the basic unit of perception is not an atomistic note irrelevant to others, but a conscious experience-fragment with a temporal depth. Each experience-fragment is composed of some number of consecutive notes around that time point, so the successive experiences are different, but are interrelated due to their overlap; (middle) A special operation at a higher conscious level is necessary to differentiate the short experience-fragments from each other, and simultaneously coalesce them in the temporal order, (right) so that the person can actually perceive the temporality or enjoy the overall beauty as well as the detailed rhythm of the melody. Such an operation, capable of revealing the past, present and future phases of a temporal object while itself surpassing the chronological order, is postulated as an inner logical structure of consciousness (e.g. the tripartite form of retention-primal impression-protention proposed by Husserl). More detailed discussions can be found in Note S1.

**Figure supplement 2.**
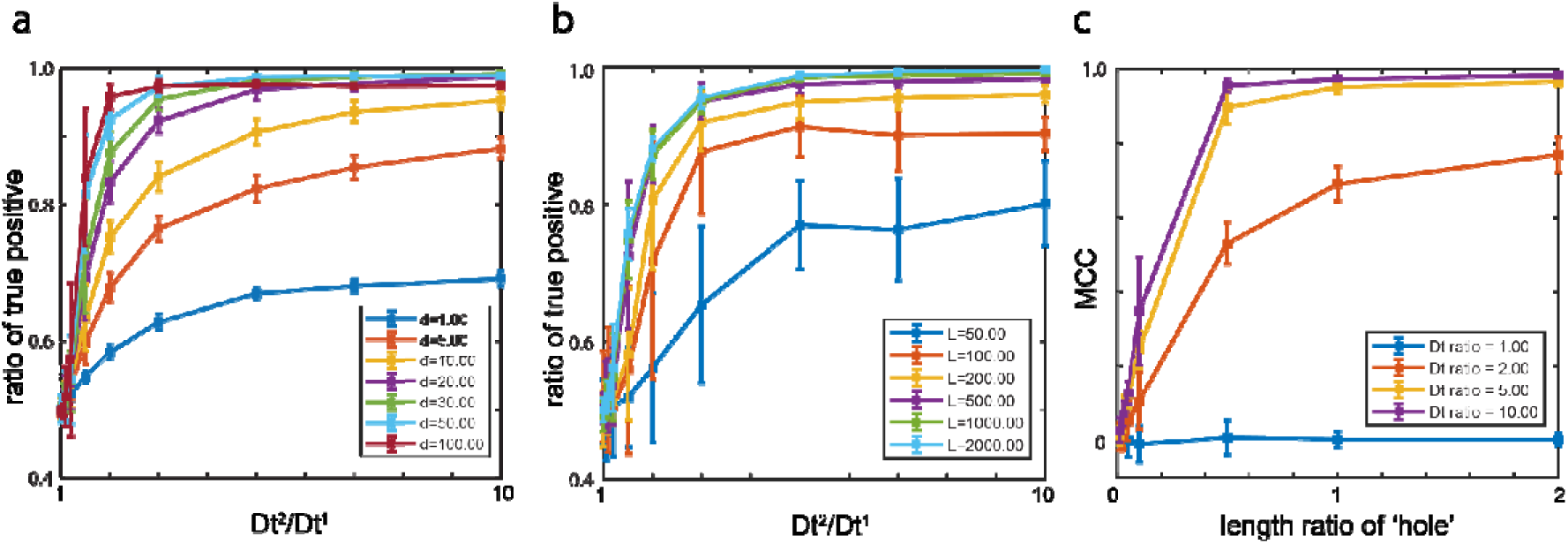
Performance validation of the SEES operator. Brownian random walk segments with different diffusion coefficient (D_t_) were merged together to evaluate the accuracy of SEES operator. All measurements were repeated 30 times, the error bar shows standard deviation of the results. (a) Two equal length of the segments with different ratio of Dt were tested under different historical experience vector length d. The differentiation accuracy was measured by the ratio of true position. Low experience vector length (d=1, blue) shows relative low accuracy even in high Dt ratio, while d=30∼50 show the best performance. Long experience vector length (d=100, dark red) shows slightly lower accuracy. (b) The same test as in a with fixed d=30 and different segment length. Short segment length (L=50, blue) shows unstable results and relative low accuracy. After the segment length reaches to 500 frames, the accuracy become satisfactory with D_t_^2^/ D_t_^1^≥3. (c) small segment with higher diffusion coefficient was inserted in a homogeneous random walk segments. The differentiation accuracy was measured by Matthews Correlation Coefficient (MCC) to address the unbalance binary classification issue, in which the fraction of correction measurement can be missleading^5^. Random results give approximately zero value of MCC as the performance in D_t_ratio = 1 group (blue), which means the diffusion coefficient of the inserted segment is equal to that of the original segments. D_t_ ratio higher than 5 and length ratio higher than 0.5 gives satisfactory results (MCC ∼ 1).

**Figure supplement 3.**
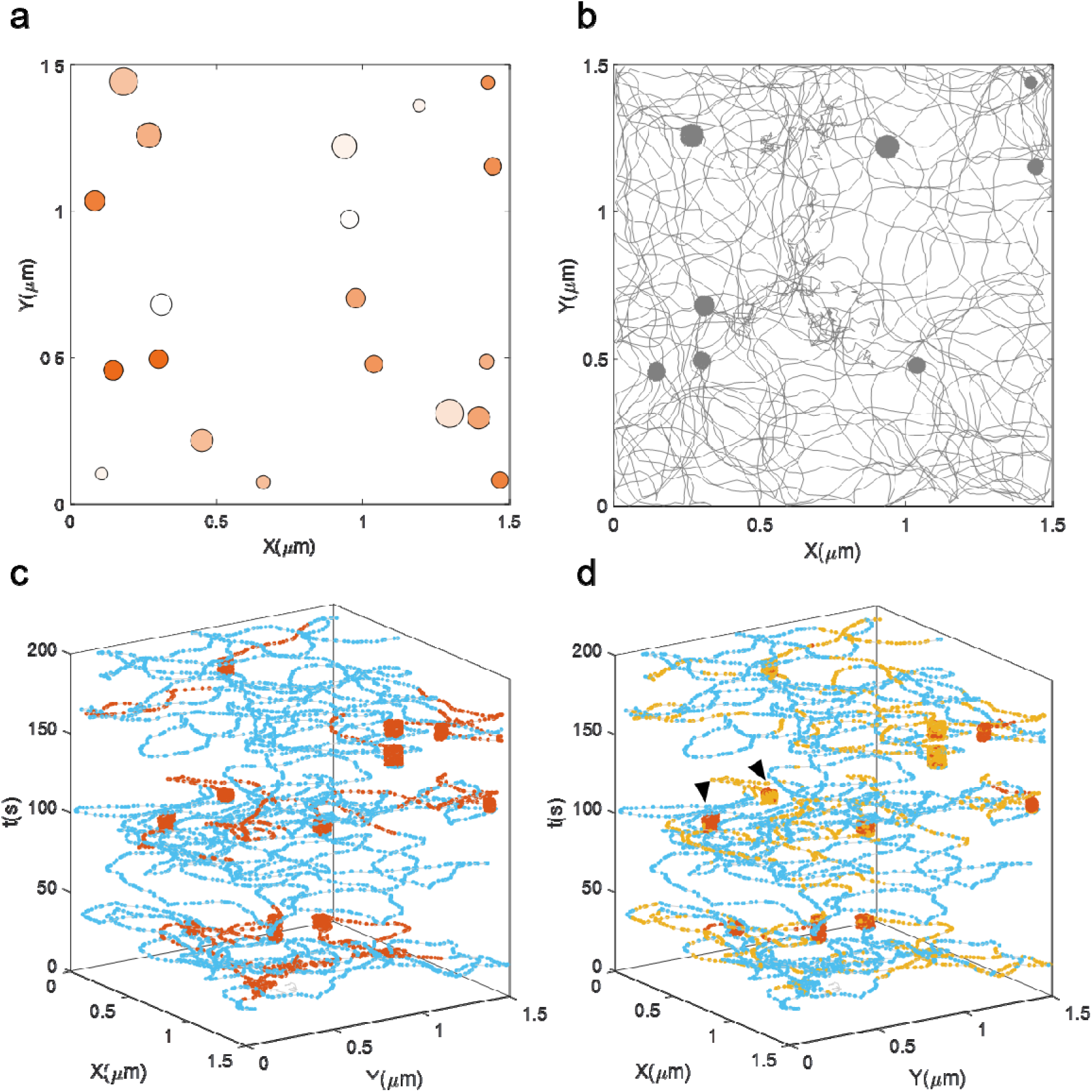
Different SEES operators reveal various underlying mechanism. (a) The simulated ground truth of the 2D cell membrane in Figure 6A. The circles represent the binding site on the membrane. Different color indicates the variations of the local diffusion coefficient of each binding site. The dark orange stands for small diffusion coefficient while the light orange represents large diffusion coefficient. (b) The simulated trajectory. (c) SEES/MSD-ED operation differentiates the binding state from the directed state. (d) SEES/U-ED operation reveals the local diffusion coefficient differences among binding sites (indicated by black arrow).

**Figure supplement 4.**
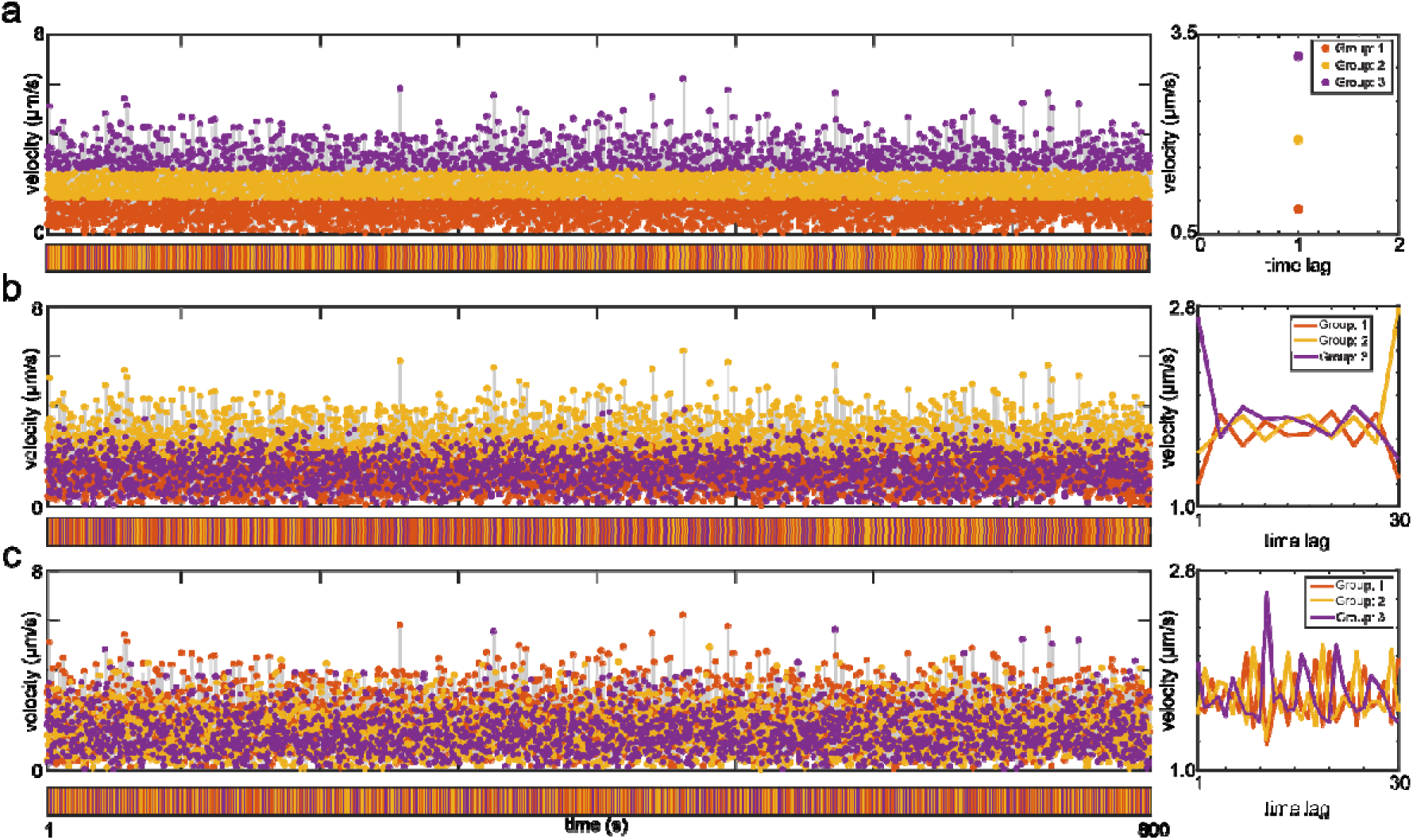
SEES operation on pure homogeneous random-walk time-series. The same parameters were used as those applied in Figure 6. Instead of assembling locally and forming colorful multiple segments as what happen in Figure 6, the same color points randomly distributed along time, regardless of the length of the experience-vectors. Furthermore, the cluster centers shown on the right also imply that there is no certain patterns found. Therefore, the collective pattern emerge with SEES operation can only originate in the underlying mechanism of the time-series data themselves but not from the clustering algorithm.

**Figure supplement 5.**
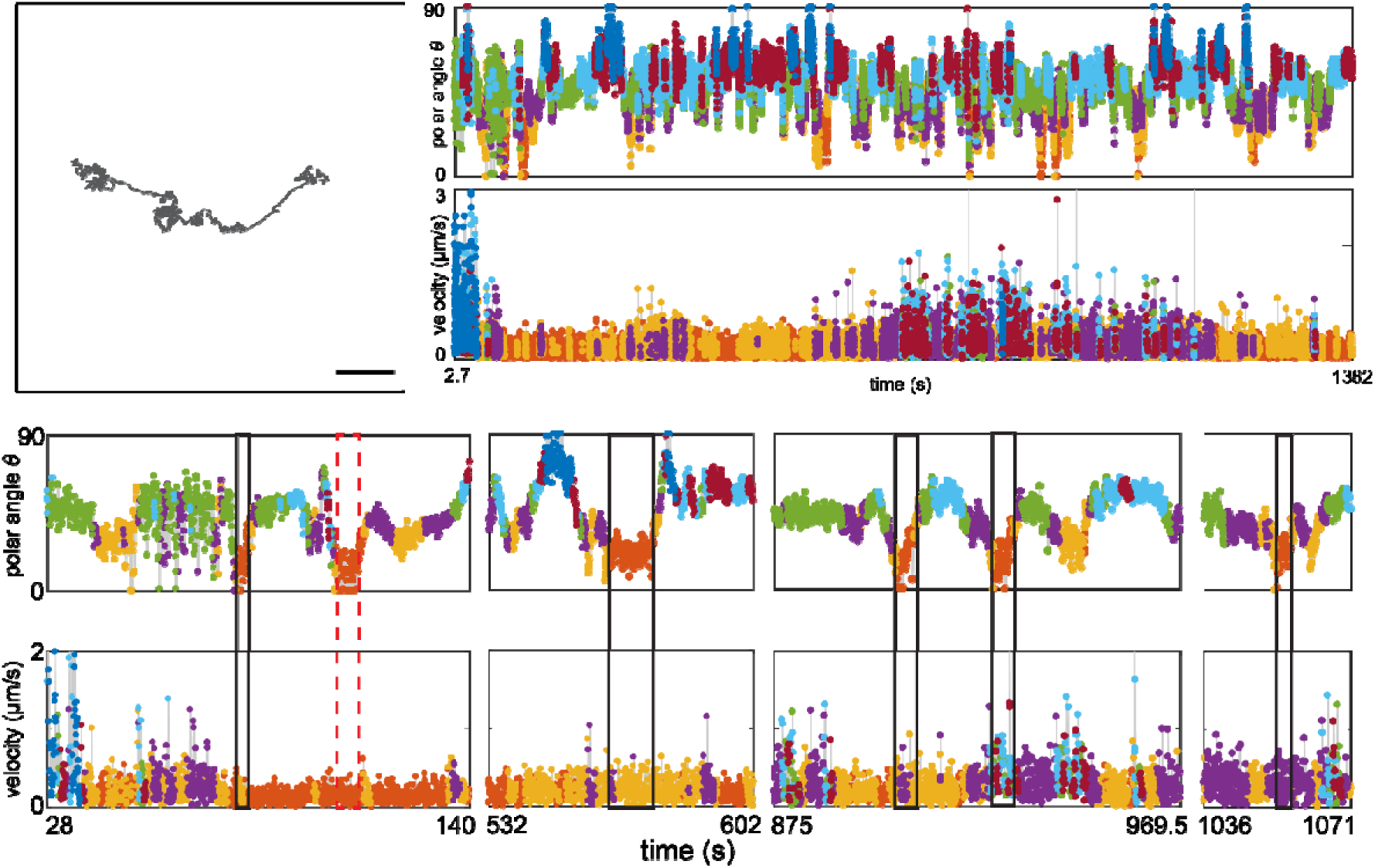
SEES/U-ED operation for cell-entry point determination of the transmembrane process of another RGD-coated AuNR. (a) The untagged trajectory. The scale bar is 3 μm. (b) The color tagged time-series and collective pattern of polar-angle (top) and velocity (bottom) variation. (c) As described in the manuscript for Figure 9B, the cell-entry-point was identified by the overlapping of same orange color in the polar-angle series and velocity series, implying the co-localization of the highest confinement, vertical particle orientation and lowest motion speed. The one with the red-dash-box shows the only perfect occurrence of this synchronization. Others are similar but failed trial sites.

**Figure supplement 6.**
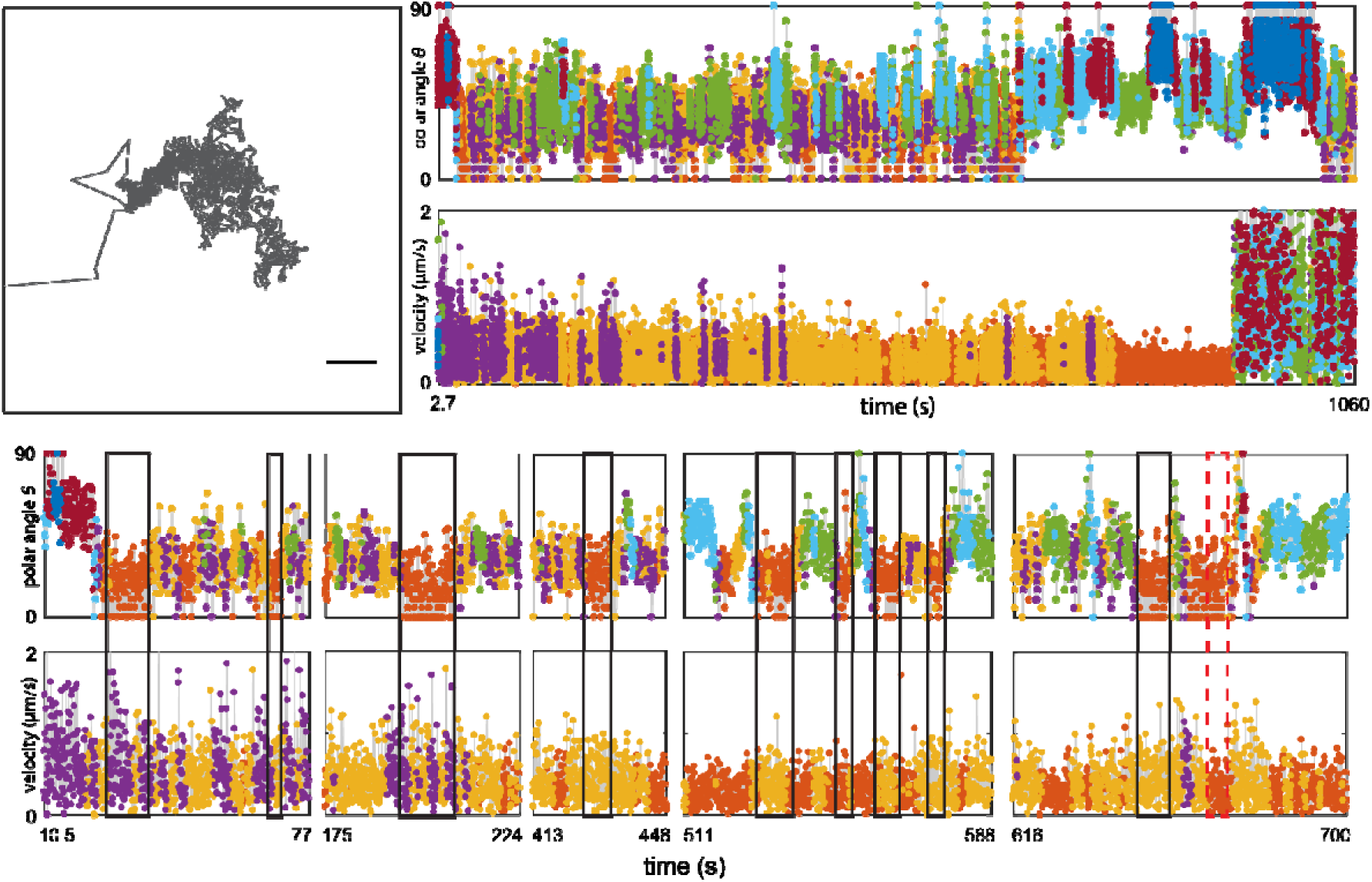
SEES/U-ED operation for cell-entry point determination of the transmembrane process of a cationic peptide coated AuNR. The scale bar is 1 μm.

**Figure supplement 7.**
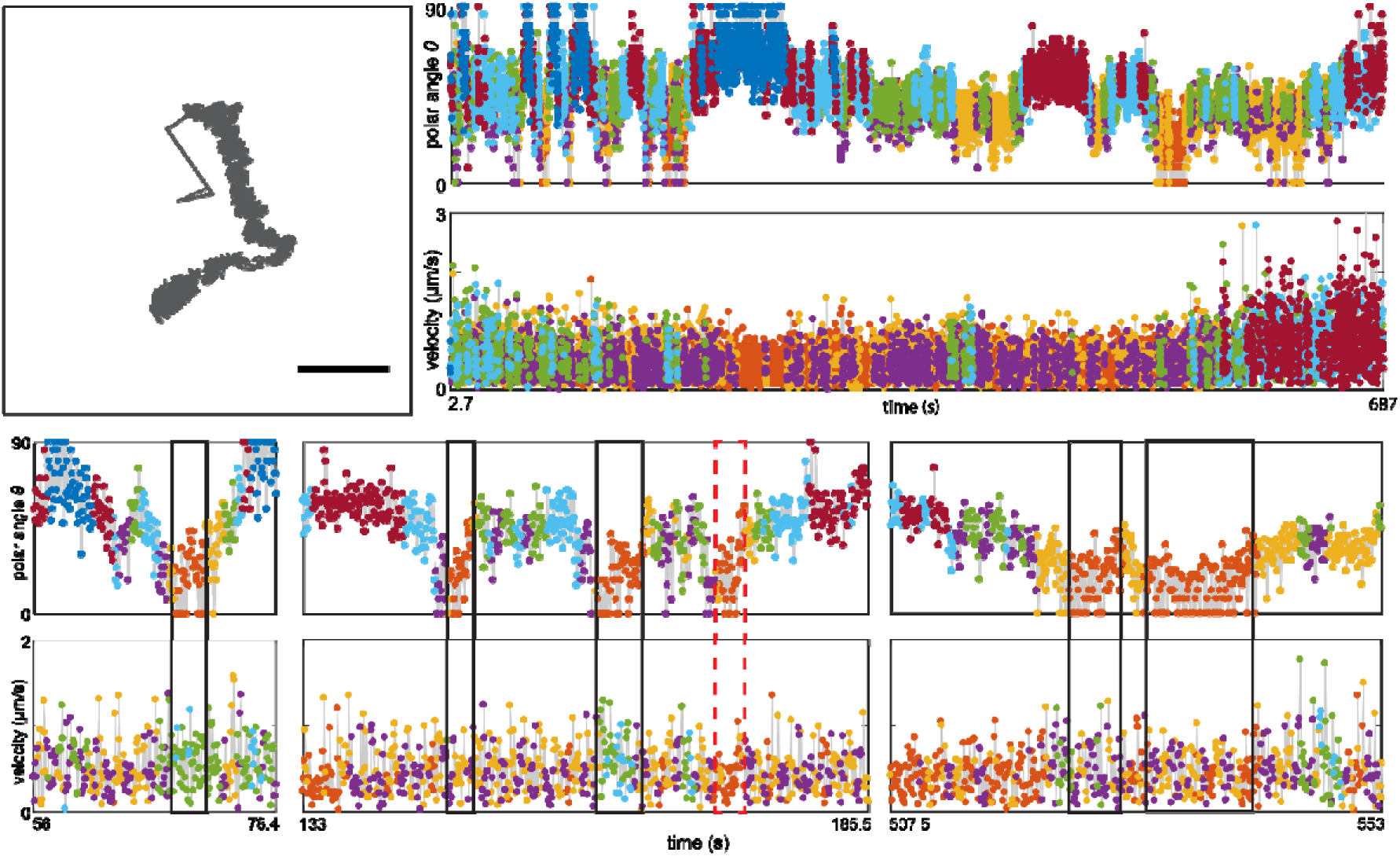
SEES/U-ED operation for cell-entry point determination of the transmembrane process of another cationic peptide coated AuNR. The scale bar is 1 μm.

## Appendix 2: responses to reviewers from the previous submission

### Reveiwer 1

In this work, the authors present a methodology for analyzing single-nanoparticle (NP) tracking data in a way that preserves spatiotemporal heterogeneity that can be lost during other analysis techniques. The method (designated by the acronym SEES) utilizes unsupervised learning (k-means clustering) to tease out dynamic differences in particle trajectories without presupposing much about the mechanisms and/or physical states of the system. In essence, the SEES technique consists of creating “overlapped historical vectors” constructed from the current data point and some number of prior data points from the particle trajectory. Next, k-means clustering is used to cluster the vectors where the number of categories (k) is determined assuming that cluster centers are local density maxima. The clusters are then projected back onto the particle time-series as different colors for visualization, thereby allowing local and global changes to become obvious as patterns in the color-coding. As a demonstration, the SEES method is used on both uni- and multi-variable simulations of single particle data. The analysis is able to resolve most of the fundamental dynamics of a rotating/revolving nanorod, as well as 1D and 2D diffusion on a membrane. However, SEES analysis of a random walk resulted in random coloring of the NP trajectory. Beyond simulation results, experimental data were collected by observing the cellular internalization process of functionalized gold nanorods (AuNRs) interacting with live (HeLa) cell membranes using darkfield microscopy. Taking advantage of the fact that the scattering color of the AuNRs reports on the spatial orientation of the particle, the authors implemented an experimental set-up that allows for simultaneous rotational and translational tracking of single AuNRs. Using SEES-analyzed trajectories of rotation angle, velocity, and position, the analysis reported on (among other things) location of the transmembrane sites, duration of cell entry, characterization of the diffusion states and transitions of the AuNR on the cell surface. Finally, SEES was used to expose the differences in transmembrane dynamics of AuNR functionalized with different RDGs and catatonic peptides, which have different cellular uptake mechanisms.

Overall, the manuscript reports a potentially useful methodology for analyzing single NP time-series data without presupposing much (if anything) about the system. This method could be useful when little about the underlying dynamics is known, or possibly as way to validate physical models by checking their assumptions against the SEES prediction. However, the basic framework behind the analytical methods are already well-known in the field (k-means clustering, for example), so the actual novelty is not immediately clear. Second, although interesting, the philosophical motivations are messy, distract from the overall message, and need to be replaced/augmented more tangible scientific arguments for clarity. In brief, the so-called “modeless” approach is motivated via an analogy to phenomenology from the field of philosophy using the example of consciousness of music. However, most of the analogy to phenomenology is unnecessary and distracts from the overall message of the paper. Perhaps it could be argued that the philosophy behind the “modeless” approach is relevant, it is largely already implied by the use of unsupervised learning.

For these reasons, and the concerns outlined in the comments below, the paper needs to be majorly revised before proceeding any further.

Reply: We thank this reviewer for the detailed summery of our work. However, we cannot agree with the conclusion of this reviewer, who belittled our model-free concept of SEES to just the unsupervised learning (k-means clustering). Indeed, while the 3 separated methods used in the SEES operator, i.e. vector construction via subseqeuncing, k-means clustering, segmenting the trajectory via coloring back, respectively, are simple and well-known, their combined use for single particle tracking (SPT) analysis has never been reported. The key novelty is the introduction of “historical analysis” into SPT analysis, i.e. to distinguish the single particle state based on its history without applying any predefined models. The unsupervised learning is only a tool to classify the historical vectors. However, to assign the momentary point on the trajectory a short history for classification and then to determine the state of the particle based on the spatial distribution of the classified points have never been proposed before. The logical foundation of this seemingly ordinary but never been thought of combination of the 3 simple operations is the philosophical arguments.

Generally, a single particle trajectory is the consequence of a free micro/nano-particle stochastically exploring an unknown complex environment for certain amount of time under optical microscopy. As a special subset of time series data, single particle trajectories have several unique characteristics. 1) Each data point results from an instantaneous reaction with the local environment and therefore has some physical meaning. 2) The reactive property of the local environment has certain continuity and the spatial and temporal scale of the continuity is larger than the spatial and temporal resolution of the optical microscopic measurement (otherwise the microscopic technique is not applicable to the particular dynamic system). Each of such a continuity could be designated as an environmental state. 3) There is an overall picture or a general story about different states of the local environment and the relationship of these states embedded in the trajectory of the single particle (otherwise no microscopic measurement or data analysis would be performed on the system). 4) Such a picture shall emerge after acquiring a sufficient but limited number of data points. Hence, the fundamental issue of SPT analysis is how to extract the embedded picture, carrying both the unity and details, from a set of discrete data points of restricted spatial and temporal range.

Previous SPT analysis are all model based, otherwise there seems no way to start with. Facing the complex situation and the awkwardness of using the known to constrain the unknown, however, we want to postpone the application of physical models as late as possible. Since there is no theory in physical science that could proceed without models, we borrowed the concept of historical analysis from social science and resort to the philosophy of phenomenology for theoretical support.

The initial motivation of this work was just to try the historical analysis approach for SPT analysis. In social life, as everyone knows, one should judge a person based on his/her historical behaviors for certain amount of time rather than his/her instantaneous picture. Therefore, when analyzing a single particle trajectory, if we use the “history” of the particle to label its dynamic state, we may avoid the too early application of physical models to oversimplify the complex system. This new method, which we eventually name it the 3-step SEES operator, returns remarkably good results in correctly identifying the NP transmembrane spot and providing more dynamic details, etc. However, when searching for the literature, we were surprised to find that there are no theoretical support on such “historical analysis” at the single particle/molecule level, and no reasonable explanation on the 3-step operation is available. Only after reading books and articles on philosophy, we realized that the so-called “history”, strictly speaking, is a phenomenological concept and would not exist without subjective interpretation (all history is contemporary history). In the fundamental research of natural science, however, there is a persistent belief that people are facing a totally objective world, and the subjective factors should be eliminated as much as possible. Therefore, an individual particle, at any moment, would have no history. Its behavior is always bound by some essential physical laws and reversible in time if all the limiting internal properties and environmental conditions are obtained. A timeless physical model or equation on the dynamic states of the particle, even though the time may be a necessary parameter, is absolutely required before analyzing its trajectory.

As natural scientists, we certainly agree with the objectiveness of our research objects, and the necessity of physical models to describe the objects. What we want to argue in this work is that it may not be necessary to apply models at the very beginning of the analysis when there is little clue on the complex system. Since data analysis is not straight forward (otherwise we would not have to analyze the data) and inevitably carries subjective factors, we have to be cautious when applying models. In order to avoid bias or oversimplification due to subjective judgement as much as possible, however, we would have to resort to our intuition. Because, according to the phenomenology study, the intuition, though transcendental, can provide an overall picture on the object and is less biased than the model-based judgement. Alternatively, if an object does carry some meaningful information or certain unity, it should be able to have the object exhibit the “truth” on its own in an unbiased way graspable by the intuition. For example, although it is possible to write down any combination of musical notes, it is a meaningful melody only when the combination, through the human sensory, passes the screening of the intuition and be interpreted by the consciousness as a melody. Only after it is recognized as a beautiful music, it becomes necessary for us to further analyze how the melody is composed based on previous musical knowledge or models. Therefore, that the SEES operator does is essentially to provide a repeatable mechanical procedure that allows the single particle trajectory to exhibit its inner unity in a “model-free” way and be graspable by the intuition, so that to facilitate further model-based analysis.

Although the unsupervised learning (k-means clustering here) allows our “model-free” approach to be implemented by classifying the input vectors without any presumptions, it does not care how the input vectors are obtained and how the classified vectors are used. In this regard, the unsupervised learning itself does not provide any solution, either conceptually or practically, to the SPT analysis problem. In this work, the assignment of a history to the otherwise atomic points, which provides inputs to the unsupervised learning, and the distribution of the same color points, which ensures the unsupervised learning results are useful, are more important than the unsupervised clustering action itself. All these are possible only with the philosophical arguments.

Taken together, the central issue of this work is on the viewpoint to the single particle trajectory. From the perspective of subject-object unification, we can assign a history to the momentary points and use their history to tell them apart and to determine their physical states. From the perspective of subject-object separation, the momentary points would have no history and can only be framed into predefined models. For this reason, the SEES operator and the model-free approach is a major breakthrough for SPT analysis.

Major comments:

1. Building on the comments above, the authors need to rewrite the manuscript to deemphasize the philosophical motivations behind the work. As only one example, the writing on this can be very unclear, for example page 5 lines 21-26.

Reply: That part of philosophical argument mentioned by this reviewer is modeled after the dialectic in Hegel’s *Phenomenology of Spirit*, which is not easy to understand for people without relevant philosophy background. We have transferred it from the main text to the Supporting Information.

2. The analogy to the gestalt of conscious of musical experience is not enough to motivate the “overlapping historical vector”. Although perhaps useful, the theoretical reasons for such a construction should be discussed and compared to other segmentation methods used in time-series clustering. What effects does the vector construction have on the properties of the data? Why does the dependence of the coloring pattern on the vector length saturate at some value?

Reply: The overlapping of historical vectors is the result of assigning each point a short history. Since it is unimaginable that an objective data point could have a “history”, as discussed earlier, the theoretical reasons cannot be found in the framework of reductionism that dominates the current scientific research. The central issue is how to understand “time”. Our thoughts are inspired by the “negation of negation of now (this moment)” elaborated by Hegel in the first chapter of his book *Phenomenology of Spirit* (Sensuous-Certainty; or the “This” and Meaning Something). Briefly, without *a priori* definitions, the “now” can only be defined by what are not the “now”, that is, to characterize the “now” point with its adjacent historical points. The construction of historical vectors happens to take the same form of subsequence or moving window operation. Since Hegel’s arguments are difficult to understand, we instead use the listening to a melody and the gestalt of musical experience as an analogy, which is also the favorite example used by Husserl to explain the structures of consciousness that make possible the experience of time. The readers are referred to the article “Phenomenology and Time-Consciousness” by Kelly (ref. 22).

Therefore, compared to the segmentation methods based on the supervised learning and the hidden Markov model (HMM), the key difference between the conventional ones and our SEES method are how to “see_”_ the data as well as the underlying states. The former ones assume a trajectory has finite number of deterministic underlying states; ours holds that the underlying states are related to the particle history and not necessarily limited in number, making it possible to discover unusual states or events. Note that neither the conventional methods nor ours can provide the true physics on the dynamic states, whose interpretation has to be derived from previous knowledge on the particular study. However, the interpretation is applied at the stage of feature selection in supervised learning or setting up the emission matrix in HMM; in our method, the interpretation is postponed to after the emergence of an intuitive pattern.

The property of the data determines which vector construction is to be used. For example, the univariate vector construction is probably more suitable for data with obvious changes in magnitude, and the autocorrelation vector construction is probably better for data with certain self-similarity. The same set of data could be readily tried with different vector construction methods, but the final choice depends on the properties of the particular system to be studied.

The convergence of the coloring pattern is one major criteria of the emergence of certain meaningful inner differences. The underlying assumption of the single particle trajectory data is that there exist temporally continuous state changes along the whole trajectory, and the particle can be retained in a state for certain period of time. A trajectory with no underlying state change such as pure random walk would never reach a stable coloring pattern with the increasing of vector length. For our evaluation or experiment data, the increasing of vector length enhances the resistance to the data noise. While the vector length is long enough to capture the historical particle dynamics, the temporal coloring pattern should converge to the underlying particle dynamics variation.

3. In the manuscript, the authors claim that the “conventional approach” of analyzing similar single particle tracking data is “inefficient and often irreproducible.” This should be demonstrated, if not by conventionally analyzing your own data as a control, then pointing to clear instances of irreproducibility in the literature.

Reply: The inefficiency and irreproducibility of SPT analysis using conventional methods mainly refers to the manual segmentation process. Since every single particle trajectory is unique, and the manual segmentation process is time-consuming, it is common for most people to pick only a few seemingly “good” trajectories from their experimental data, and analyze them based on personal experience. But such a rule-of-thumb process would not be acknowledged explicitly in any literature.

4. Similar to point 3, for the simulation of the 2D cell membrane-nanoparticle interaction --what happens if one assumes a two-state model and fit the data? Does it look like a noisy two-state system or is the 2-state assumption self-evidently wrong from the fit? The SEES method could be useful as a protection against over-simplified models.

Reply: For the simulation of the 2D cell membrane-nanoparticle interaction, if only 2 clusters are used, there is still a similar pattern, but the difference in the NP binding affinities in various confined regions will not be clearly evident.

5. The physical meaning of the AuNR SEES results could be analyzed more. How does the duration of membrane crossing compare to the literature values? Does the analysis elucidate some unique dynamic missed by and/or incongruous with conventional analysis?

Reply: The main purpose of this work is to introduce the SEES method. The applications of the method on the transmembrane process of single AuNRs are just to demonstrate that large amount of information could be obtained. For the same trajectory, we have shown that the conventional manual segmentation method returns much smaller number of fragments than the SEES method. The in-depth analysis of the physical meaning of these or similar results will be left to further studies. Just as an example, from the sequential evolution of experience-vectors (Figure 10 and Movie 6), we have identified the hop-and-linger behavior of the AuNR as it searches for the cell-entry site in its first diffusive region. We have also compared the diffusion dynamics of AuNRs with different surface modification, which have not been reported before. Of course, people may argue that the same information could also be obtained with conventional SPT analysis methods. However, the SEES method is not an incremental modification of an established method, therefore, other than pointing out the clear difference in the theoretical foundation, we think it is not appropriate to compare the analysis results on any particular single particle trajectory. Different researchers obtain different experimental data. Any method could be a good method as long as it serves the purpose.

6. The authors mention that subsequence clustering of time-series similar to the method used in this manuscript can result in physically meaningless clusters. How does one differentiate between a result that has no physical interpretation and one that is simply been missed so far by conventional techniques that collapse spatiotemporal heterogeneity? Additionally, it has been proposed that subsequence clustering of time series results in cluster centers that are essentially random.

Reply: In the revised manuscript, we emphasize that our method serves as a preprocessing technique to give an intuitive picture of the complicated SPT data. Over-simplification and over-interpretation can be on the two opposite sides of the data mining. Our method does not guarantee that all the emerging patterns have certain physical meaning, as do all unsupervised learning methods. It just provides clues for further research with less information loss. Though the conventional methods give physical meaning before the data analysis based on prior knowledge, try-and-error and manual post-interpretation are usually involved in the workflow. To confirm that a new dynamic pattern is found by SEES, there are a few general steps. First, ensure the emerging pattern is stable with slight experience vector length variation or different way of vector construction. Secondly, compare the obtained colored trajectory with the expected physics or dynamics, and determine whether the pattern is a reasonable particle-environment interaction. Third, analyze the pattern by conventional physical models to get important parameter values on the dynamics, such as diffusion coefficient, velocity, etc. Finally, come up with new hypothesis about the underlying mechanism and confirm it with previous publications or further experiment. We note that our method is not universal to address all the problems during data mining, but just provides valuable clues for the researchers to see their SPT data with minimal information loss.

7. How does noisy data affect the SEES result? It would be interesting to compare data collected with the focal plane of the microscope at the cell top with data taken at the cell side. The former will have more noise due to increased Rayleigh scattering from the intercellular structures since it is at the thickest part of the cell. This interferes with the plasmonic measurement.

Reply: By gradually increasing the length of the experience vector, the SEES method can resist the random noise to some extent and converge to a stable color pattern. We have performed simulations on the noise effect and obtained satisfactory results. Under the real experimental situations, however, it is not appropriate to do the comparison since no two single particle trajectories are the same. Again, the SEES method is only a preprocessing approach, and the obtained patterns need to be evaluated according to previous knowledge.

### Reviewer: 2

The paper is reasonably well organized, although it should be edited for English. The strategy proposed is novel compared to standard physics based trajectory analysis in colloid science. There are a number of comments and questions for the authors:

1. The approach appears to lack any physical basis or theoretical foundation but appears to be more of an ad hoc technique. For example, the choice of vector length requires sufficient tests when ground truths are available. When the ground truth is not available, the choice of experience vector length also requires assumptions.

Reply: The target our method faces is those biophysical systems which are lack of fully descriptive physical models. In this case, our method enables the researchers have a first glance of the obtained data with as minimal information losses as possible. The pre-suppositionless method proposed here achieves this goal by constructing experience vectors without pre-defined model based feature extraction and coloring the trajectories with the emerged inner differences.

As an exploration method, the SEES operation has no “ground truth” in real-world application. Two criteria can be adopted here to infer whether the algorithm gives a reasonable result. First, self-organized small fragments with the same color should appear along the trajectories based on the continuity underlying states change assumption. Total random series with no meaningful information were colored randomly with any vector length as shown below.

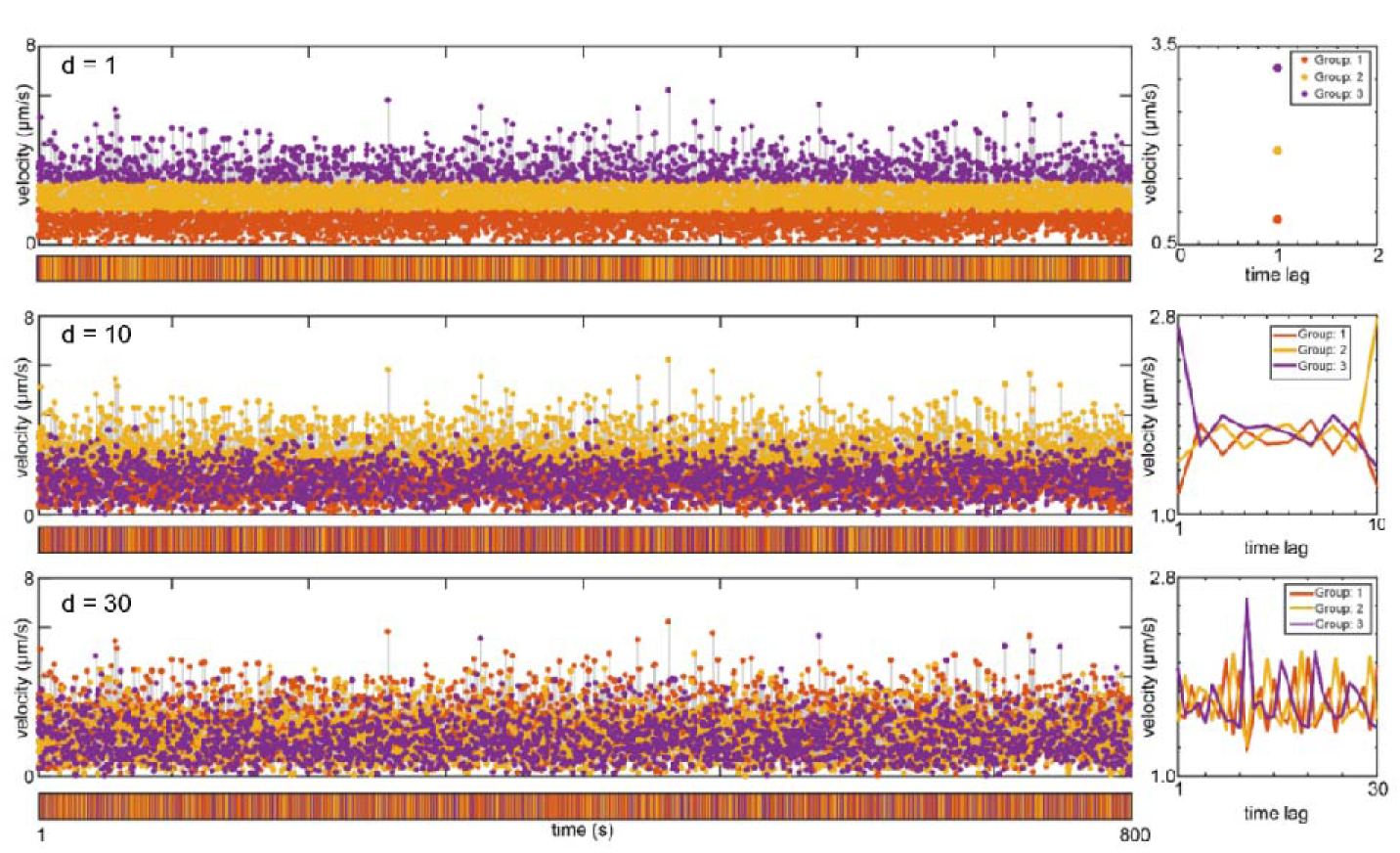

Secondly, the coloring fragment distribution should converge to a stable pattern with increasing vector length, which indicates that the experience vector is long enough to capture the historical particle dynamics and robust to detection noise.

2. Given the wide availability of machine learning algorithm libraries (e.g., Python, R, Matlab etc.) and measurement data, one of the major challenges in machine learning problems is feature engineering (i.e., the input to the machine learning algorithm). Although the authors used experience vector as a feature vector and obtain satisfactory results, there is little discussion on how to select features in different dynamic problems and how to select distance metrics between features. From this perspective, the method is very likely to see only limited use to the analysis of NP dynamics in well-understood scenarios.

Reply: As this reviewer indicates, in the proposed method, we use the experience vector instead of feature vector to describe the particle dynamics. The reason for this design is to eliminate the information loss as much as possible introduced by pre-defined model/statistics based raw data fitting/averaging.

In our method, the way of experience vector construction (univariate, MSD, autocorrelation) and the choices of the metrics for comparison reflect how the researchers want the data being interpreted. Briefly, vector construction using Univariate focuses on the overall tendency, MSD focuses on the confinement in state space, and Auto-Correlation focuses on the self-similarity of the historical dynamics. As for comparison metrics. Euclidean distance is more sensitive to the magnitude of the vector value/magnitude while the Cross-Correlation distance is more sensitive to the tendency.

For a specific single particle trajectory, the choice of the algorithm setup depends on how the researchers want to ‘see’ the data. For example, to see magnitude variation pattern along the series, the Univariate construction and Euclidean metric are a good choice. To see the confinement states, MSD construction and Cross-Correlation metric are the first choice.

Besides, different choices of the experience vector construction methods (Univariate & MSD) help to interpret the simulated data in different perspectives to reveal more detailed information of the particle dynamics (local viscosity & bonding state). To our knowledge, the compositions of these experience vector construction methods and the vector comparison methods have covered the most situations in SPT analysis.

3. The reason investigators use models to classify and analyze dynamic trajectories is for better interpretation of the underlying physical process. The model-free approach can be viewed as a useful data pre-processing step; however, a model is still ultimately needed to interpret underlying physics.

Reply: Our method indeed aims at serving as pre-processing tool to give an intuitive picture of the complex particle dynamics with as less predefined model as possible. Manual interpretation, assumptions and physical models are still essential for the researchers in the further investigation steps. The novelty and significance of our manuscript is to provide with a concept and robust method that put the pre-suppositionless data mining and visualization step at the very beginning instead of at the end of the experiment data investigation. While the predefined model may cause oversimplification or information loss by fitting or averaging, our method lets the inner differences in the data emerge themselves to support the following investigation and modeling, which is of special significance when dealing with previously poorly explored systems such as AuNR-cell membrane interactions. Overall, by providing an objective, information retained picture of the system for further investigation, our method can be a complementary pre-viewing steps for the model-based workflow.

4. The test cases considered in this paper is not sufficiently complex to demonstrate the effectiveness of this method compared to other methods cited by the authors. The authors should benchmark their approach against model-based approaches in a complicated scenario to demonstrate the usefulness of their approach.

Reply: Following this reviewer’s advice, we applied our method to the evaluation data of a recent published paper (Nat. Commun. 2015). In this paper, a persistence-active model was proposed to analyze the cell migration trajectory. Without knowing the underlying model and mechanism, our method provides reasonable coloring pattern comparing to the ground truth and successfully reveals the underlying dynamics in the evaluation data while no prior-knowledge on the system is involved. As the underlying mechanism of cell migration is largely different from that of the NP-cell interaction, the results indicates that our method is reliable and potentially applicable for wide research field based on single particle tracking.

5. This method proposed is not entirely new. The idea of using a sequence as input is quite common in machine learning for natural language processing and in signal processing to classify time series. For example, in natural language processing, a recurrent neural network is used to cluster sequences with automatically extracted features.

Reply: Our SEES method consists of three main steps: experience vector construction, unsupervised comparison and trajectory coloring. For each step, the underlying algorithm indeed previously exists. However, putting them all together, and establishing the pre-suppositionless SPT analysis concept and the operation pipeline are for the first reported in this manuscript to the best of our knowledge.

Comparing to the recurrent neural network (RNN) in natural language processing, which extracts local and long-term features from the input sequence by iteratively minimizing loss function, in our method numbers of experience vectors are constructed from the input sequence with deterministic steps. Besides, while comparison can potentially be a task of RNN (classification task), it’s essential for the second step in SEES method.

SEES method distinguishes from the neural network based methods by its ability of operating without a specific objective function. As a data exploration tools, it gives an information preserved intuitive picture of the system. Meanwhile, as a pre-suppositionless method, SEES method is also different from pre-defined model based method for its capability of dealing with various single particle time series observations of totally different physical meaning (such as velocity and polar angle) without much modification.

